# Modeling behaviorally relevant neural dynamics enabled by preferential subspace identification (PSID)

**DOI:** 10.1101/808154

**Authors:** Omid G. Sani, Bijan Pesaran, Maryam M. Shanechi

## Abstract

Neural activity exhibits dynamics that in addition to a behavior of interest also relate to other brain functions or internal states. Understanding how neural dynamics explain behavior requires dissociating behaviorally relevant and irrelevant dynamics, which is not achieved with current neural dynamic models as they are learned without considering behavior. We develop a novel preferential subspace identification (PSID) algorithm that models neural activity while dissociating and prioritizing its behaviorally relevant dynamics. Applying PSID to large-scale neural activity in two monkeys performing naturalistic 3D reach-and-grasps uncovered new features for neural dynamics. First, PSID revealed the behaviorally relevant dynamics to be markedly lower-dimensional than otherwise implied. Second, PSID discovered distinct rotational dynamics that were more predictive of behavior. Finally, PSID more accurately learned the behaviorally relevant dynamics for each joint and recording channel. PSID provides a general new tool to reveal behaviorally relevant neural dynamics that can otherwise go unnoticed.

## Introduction

Modeling of how behavior is encoded in the dynamics of neural activity over time is a central challenge in neuroscience. This modeling is essential for investigating or decoding behaviorally measurable brain functions such as movement planning, initiation and execution^1–3^, speech and language^4^, mood^5^, decision making^6^, or neurological dysfunctions such as movement tremor^7^. However, building such models is challenging for two main reasons. First, in addition to the behavior being studied, recorded neural activity also encodes other brain functions, inputs from thousands of other neurons, as well as internal motivational states with brain-wide representations such as thirst^3,8–15^. These together constitute behaviorally irrelevant neural dynamics. Second, many natural behaviors such as unconstrained movements or speech are temporally structured. Thus understanding their neural representation is best achieved by learning a dynamic model, which explicitly characterizes the temporal evolution of neural population activity^3,16–18^. Given these two challenges, answering increasingly sought-after and fundamental questions about neural dynamics such as their dimensionality^3,13,19^ and important temporal features such as rotations^14,20–22^ requires a novel dynamic modeling framework that can prioritize extracting those neural dynamics that are related to a specific behavior of interest. This would ensure that behaviorally relevant neural dynamics are not masked or confounded by behaviorally irrelevant ones and will broadly impact the study of diverse brain functions. Developing such a dynamic modeling framework has remained elusive to date.

Currently, dynamic modeling of neural activity is largely performed according to two alternative conceptual frameworks. In the first framework, often termed representational modeling (RM), behavioral measurements such as movement kinematics, choices or tremor intensity at each time are assumed to be directly represented in the neural activity at that time^2,7,23,24^. By making this assumption, RM implicitly assumes that the dynamics of neural activity are the same as those in the behavior of interest; the RM framework thus takes behavior to represent the brain state in the model and learns its dynamics without considering the neural activity (Fig. 1a; Methods). This assumption, however, may not hold since neural activity in many cortical regions including the prefrontal^6,25^, motor^20,26–28^ and visual^13^ cortices and other brain structures such as amygdala^8,9^ is often simultaneously responsive to multiple behavioral and task parameters^6,8,9,25^ and thus is not fully explained by the RM framework^3,17,18,20^. Motivated by this complex neural response, recently a second framework known as neural dynamic modeling (NDM) has received growing attention^3,5,16,18,20–22,29–32^ and has led to recent findings for example about movement generation^3,20^ and mood^5^. In NDM, the dynamics of neural activity are modeled in terms of a latent variable that constitutes the brain state in the model and is extracted purely using the recorded neural activity and agnostic to the behavior (Fig. 1a). Once extracted, this latent brain state is then assumed to encode the behavior of interest, such as movement kinematics^21,29,30^ or mood^5^. Because NDM does not guide the extraction of neural dynamics by behavior, it may miss or less accurately learn some of the behaviorally relevant neural dynamics, which are masked or confounded by behaviorally irrelevant ones. Uncovering these behaviorally relevant neural dynamics requires a new modeling framework to extract the dynamics that are shared between the recorded neural activity and behavior of interest, rather than extracting the prominent dynamics present in one or the other as done by current dynamic models (Fig. 1a)—present in behavior in the case of RM and in neural activity in the case of NDM.

**Figure 1.**
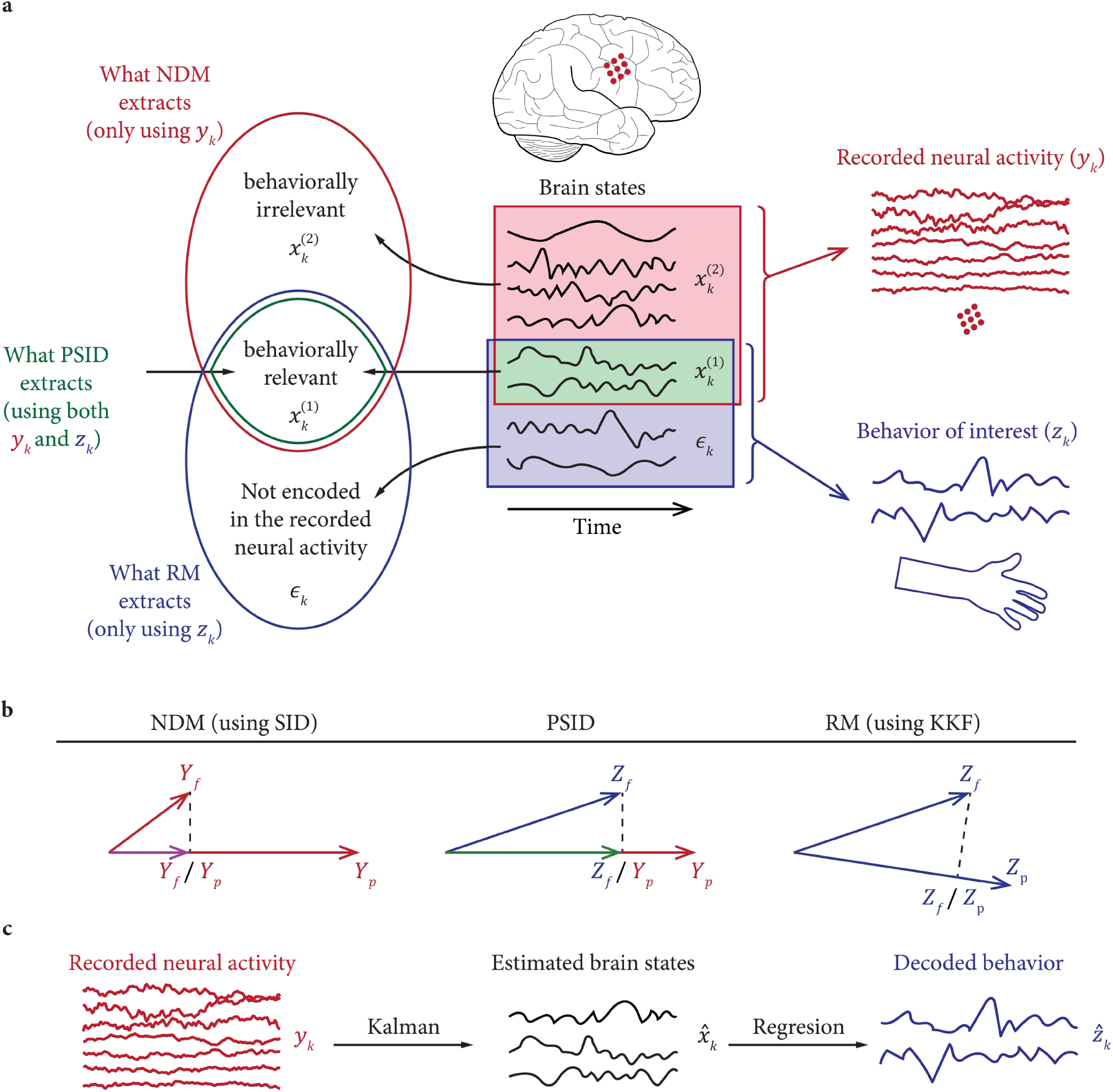
PSID enables learning of dynamics shared between recorded neural activity and measured behavior. (**a**) Schematic view of how the state of the brain can be thought of as a high-dimensional time varying variable of which some dimensions (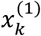 and 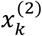) drive the recorded neural activity (*y*_*k*_), some dimensions (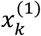 and *ϵ*_*k*_) drive the measured behavior (*z*_*k*_), and some dimensions 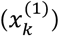 drive both and are thus shared between them. The choice of a learning method affects the brain states that are extracted from neural activity. NDM extracts states regardless of their relevance to behavior and RM extracts states regardless of their relevance to recorded neural activity. PSID enables extraction of brain states that are related to both the recorded neural activity and a specific behavior. (b) Schematics of how PSID achieves its goal in comparison with a representative NDM method (i.e. SID) and an RM method (i.e. KKF). *A*/*B* denotes projecting *A* onto *B* (Methods). The key idea in PSID is to project future behavior *z*_*k*_ (denoted by *Z*_*f*_) onto past neural activity *y*_*k*_ (denoted by *Y*_*p*_). This is unlike NDM using SID, which instead projects future neural activity (denoted by *Y*_*f*_) onto the past neural activity *Y*_*p*_ (Methods). It is also unlike RM using KKF, which projects future behavior onto past behavior (denoted by *Z*_*p*_). (**c**) For all three methods, after the model parameters are learned, the procedures for state estimation and neural decoding of behavior are the same. A Kalman filter operating on the neural activity estimates the brain states, and behavior is decoded by applying a linear regression to these estimated brain states (Methods).

In this Technical Report, we develop a novel general modeling and learning algorithm, termed preferential subspace identification (PSID), for extracting and modeling behaviorally relevant dynamics in high-dimensional neural activity. PSID uses both neural activity and behavior together to learn (i.e. identify) a dynamic model that describes neural activity in terms of latent states while prioritizing the characterization of behaviorally relevant neural dynamics. The key insight in PSID is to identify the subspace shared between high-dimensional neural activity and behavior, and then extract the latent states within this subspace and model their temporal structure and dynamics (Methods).

We first show with extensive numerical simulations that PSID learns the behaviorally relevant neural dynamics significantly more accurately, with markedly lower-dimensional latent states, and orders of magnitude fewer training samples compared with standard methods. We then demonstrate the new functionalities that PSID enables by applying it to large-scale motor cortical activity recorded in two non-human primates (NHP) performing an unconstrained naturalistic 3D reach, grasp, and return task. We show that PSID uniquely uncovers several new features of neural dynamics underlying motor behavior. First, PSID reveals that the dimension of behaviorally relevant neural dynamics is markedly lower than what standard methods conclude. Second, while both NDM and PSID find rotational neural dynamics during our unconstrained 3D task, PSID uncovers rotations that are in the opposite directions in reach vs return epochs and are significantly more predictive of behavior compared with NDM, which in contrast finds rotations in the same direction. Third, compared with NDM and RM, PSID more accurately learns behaviorally relevant neural dynamics for almost all of the 27 arm and finger joint angles and for 3D end-point kinematics. Finally, PSID reveals that almost all individual channels across the large-scale recordings have behaviorally relevant dynamics that are learned more accurately using PSID.

## Results

### Overview of PSID

We consider the state of the brain at each point in time as a high-dimensional latent variable of which some dimensions may drive the recorded neural activity, some may drive the observed behavior, and some may drive both (Fig. 1a). We thus model the recorded neural activity 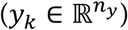 and behavior 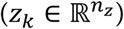 using the following general dynamic linear state-space model (SSM) formulation

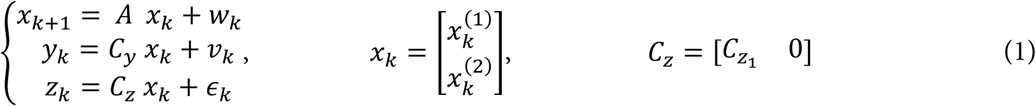

where 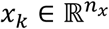 is the latent brain state that drives the recorded neural activity, and 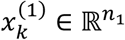 and 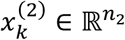 (with *n*_2_ = *n*_*x*_ − *n*_1_) are its behaviorally relevant and behaviorally irrelevant components, respectively. The matrix *C*_*Z*_ is non-zero only in its first *n*_1_ columns (i.e.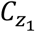) indicating that 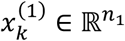 drives the behavior but 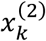 does not. Finally, *ϵ*_*k*_ represents the behavior dynamics that are not present in the recorded neural activity, and *w*_*k*_ and *v*_*k*_ are noises. *A, C*_*y*_, and *C*_*z*_ and noise statistics are model parameters to be learned using PSID given training samples from neural activity and behavior (Methods). This provides a general formulation whose special cases also include standard NDM (when *n*_2_ = 0 and *C*_*z*_ is a general matrix to be learned) and RM (when *C*_*z*_ is identity and *ϵ*_*k*_ = 0, Methods).

The goal of PSID is to build a model for how high-dimensional neural activity evolves in time while prioritizing the behaviorally relevant neural dynamics, which are the ones driven by the behaviorally relevant latent states (i.e. 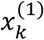, Methods). The key idea for achieving this goal is the demonstration that the behaviorally relevant latent states lie in the intersection of the space spanned by the past neural activity and the space spanned by the future behavior (Methods). Using this idea, we can extract the behaviorally relevant latent states via an orthogonal projection of future behavior onto the past neural activity (Fig. 1b, Methods). The remaining neural dynamics correspond to the latent states that do not directly drive behavior (i.e.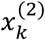). These remaining latent states can then be extracted by an additional orthogonal projection from the residual neural activity (i.e. the part not predicted by the extracted behaviorally relevant latent states) onto past neural activity (Methods). Finally, model parameters that describe the temporal evolution can be learned based on the extracted latent states. Thus, PSID solves two challenges. It builds a dynamic model of how high-dimensional neural activity evolves in time (temporal structure) and at the same time dissociates behaviorally relevant and irrelevant dynamics.

We compare PSID with standard NDM and RM. Standard NDM describes neural activity using a latent SSM that is a special case of that in PSID (equation (1)), but in terms of a latent state that is learned agnostic (i.e., unsupervised) with respect to behavior^5,21,29,30^; it then regresses the latent states onto the behavior^5,21,29^. Since standard NDM methods extract the latent states and learn their dynamics without using the observed behavior, unlike PSID, they do not prioritize the behaviorally relevant neural dynamics. While there are various methods to learn the latent SSM from neural data in the case of NDM, we use the standard subspace identification (SID) algorithm^33^, which has been used for NDM before^5,32,34^ and like PSID has a closed-form solution^33^ and is thus computationally efficient. SID identifies the latent states by projecting future *neural activity* onto past neural activity (Fig. 1b) in contrast to PSID that projects future *behavior* onto past neural activity (Fig. 1b). As control analyses, we also repeat some key NDM analyses with Expectation Maximization (EM) that can also be used to learn the model in NDM but is iterative and thus computationally complex. To implement RM^2,23^, we use the commonly-used RM method (sometimes termed Kinematic-state Kalman Filter (KKF)^21^), which builds an auto-regressive model for the behavior and directly relates the behavior to the neural activity using linear regression^2,23^. RM learns the state and its dynamics agnostic to the observed neural activity (Fig. 2b) and thus, as we will show, may learn state dynamics that are not encoded in the observed neural activity.

**Figure 2.**
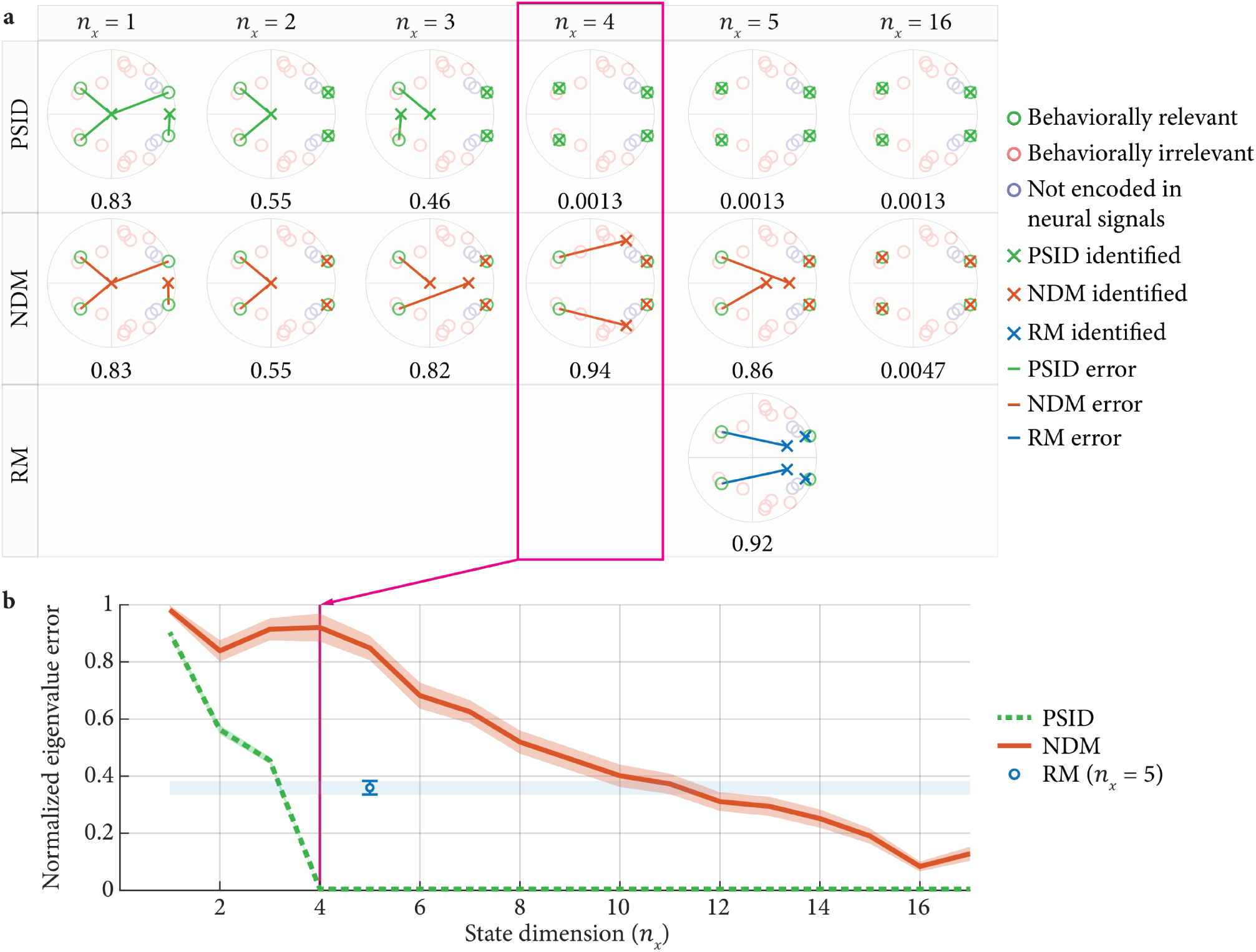
PSID correctly learns the behaviorally relevant dynamics even when using fewer latent states and performing dimensionality reduction in contrast to standard methods. (**a**) For one simulated model, the identified behaviorally relevant eigenvalues are shown for PSID, NDM, and RM and for different latent state dimensions. For RM, the state dimension can only be equal to the behavior dimension (here *n*_*z*_ = 5). Eigenvalues are shown on the complex plane, i.e. real part on the horizontal axis and imaginary part on the vertical axis. The unit circle is shown in gray. True model eigenvalues are shown as lightly colored circles, with colors indicating their relevance to neural activity, behavior, or both. Crosses show the identified behaviorally relevant eigenvalues. Lines indicate the identified eigenvalue error whose normalized value—average line length normalized by the average true eigenvalue magnitude is noted below each plot (Methods). (**b**) Normalized eigenvalue error given 10^6^ training samples is shown when using PSID, NDM and RM, averaged over 100 random models. For all random models, the total number of latent states (*n*_*x*_ = 16), the number of behaviorally relevant states (*n*_1_ = 4), and the number of behavior dimensions not encoded in neural activity (i.e. 4) is as in (a). Solid lines show the average error and shaded areas show the s.e.m. For NDM and PSID, total state dimension is changed from 1 to 16 (for PSID *n*_1_ = 4). Since for RM the state dimension can only be equal to the behavior dimension (*n*_*z*_ = 5), for easier comparison, the RM s.e.m is shown as error bars and also a horizontal shaded area.

Importantly, all three methods (RM, NDM, PSID) describe the neural activity using the same model structure, which is a linear SSM (Methods). The critical difference is how states and their dynamics are learned from neural data (NDM), from behavior data (RM) or from both (PSID), and thus which brain states are extracted (Fig. 1a). After SSM model parameters are learned in each of these three methods, in all of them, the estimation of the state from neural activity and the decoding of behavior are done using a Kalman filter and linear regression, respectively (Fig. 1c).

### Neural Recordings

We first validated PSID using extensive numerical simulations and then used PSID to uncover the behaviorally relevant neural dynamics in large-scale cortical recordings of two adult Rhesus macaques performing unconstrained naturalistic 3D reach, grasp, and return movements (Methods). In each trial, this task requires the monkey to reach for an object, grasp it, and then release the object and return the hand to the resting position. The angle of 27 (monkey J) or 25 (monkey C) joints on the right shoulder, elbow, wrist, and fingers at each point in time is tracked via reflective markers and is taken as the behavior of interest (Methods). In addition to joint angles, we also study the 3D end-point position of hand and elbow as the behavioral measurements. Large-scale neural activity was recorded from primary motor cortex (M1), dorsal premotor cortex (PMd), ventral premotor cortex (PMv), and prefrontal cortex (PFC) and for monkey C also included ipsilateral coverage (Methods). We used the local field potential (LFP) power in 7 frequency bands as the neural features to be modeled (Methods, Discussion). We use the cross-validated correlation coefficient (CC) of decoding behavior using neural activity as the main measure for how accurately the behaviorally relevant neural dynamics are learned.

### PSID correctly learns all the model parameters

We first performed simulations and found that the PSID algorithm can correctly identify all the true model parameters from data. We generated 100 validation models with random parameters and simulated sample data from each model (Methods). We then performed model identification with the PSID algorithm and evaluated the error in identification of all model parameters (Supplementary Fig. 1). We found that all model parameters were identified with less than 1% error (Supplementary Fig. 1a). Also, the identification error consistently decreased as the number of training samples increased, suggesting that even smaller errors can be achieved using more training samples (Supplementary Figure 1b). Also, compared with standard SID, PSID showed a similar error and rate of convergence (Supplementary Fig. 1c, d), indicating that even when learning of all latent states is of interest rather than just the behaviorally relevant ones, PSID performs as well as SID. Finally, we found that given a fixed training sample size, the identification error of both PSID and SID for different random models was significantly correlated with a mathematical measure of how inherently difficult it was to extract the latent states in these models from data (Supplementary Fig. 2); this indicates that with sufficient training data, even models that are inherently more difficult to learn can eventually be identified accurately. Together, these results show that PSID can correctly identify both the behaviorally relevant and irrelevant latent states.

In the above analysis, for each true validation model, we used PSID to fit a model with the same model structure parameters *n*_*x*_ and *n*_1_ as the true model (Methods). We next found that using a cross-validation procedure (Methods), we could accurately estimate both model structure parameters from training data (Supplementary Fig. 3). *n*_*x*_ and *n*_1_ were estimated with no error in 98% and 94% of the models, respectively; also, their average estimation errors were 0.040 ± 0.028 (mean ± s.e.m.) and 0.050 ± 0.021, respectively (Supplementary Fig. 3a, c). The error in estimating *n*_*x*_ was similar to that achieved when using the same cross-validation procedure for the standard SID (0.08 ± 0.039), which also has the parameter *n*_*x*_ (Supplementary Fig. 3b).

### PSID prioritizes identification of behaviorally relevant dynamics

We found that, unlike standard methods, PSID correctly prioritizes identification of behaviorally relevant dynamics even when performing dimensionality reduction, i.e., even when identifying models with fewer latent states than the total number of latent states in the true model. We applied PSID to simulated data from 100 random validation models with 16 latent states (*n*_*x*_ = 16) out of which 4 were behaviorally relevant (*n*_1_ = 4). We used PSID to identify models with different latent state dimensions and evaluated how closely the identified latent state dynamics matched the true behaviorally relevant latent state dynamics. As the main performance measure, we computed the identification error for learning the eigenvalues of the behaviorally relevant component of the state transition matrix *A* (Methods). These eigenvalues specify the frequency and decay rate of the response of the latent states to excitations (i.e. *w*_*k*_) and thus determine their dynamical characteristics (Methods). The location of eigenvalues in the true and identified models is illustrated in Fig. 3a for one of the validation models. We found that PSID accurately identifies the behaviorally relevant latent states while standard methods can identify latent states that are unrelated to behavior (NDM), or latent states that are not encoded in the observed neural activity (RM). Overall, using a total latent state dimension of 4, PSID learned all 4 behaviorally relevant eigenvalues while the standard methods could not (Fig. 2a); further, PSID achieved higher accuracy compared with standard methods even when they used higher dimensional latent states (Fig. 2b).

**Figure 3.**
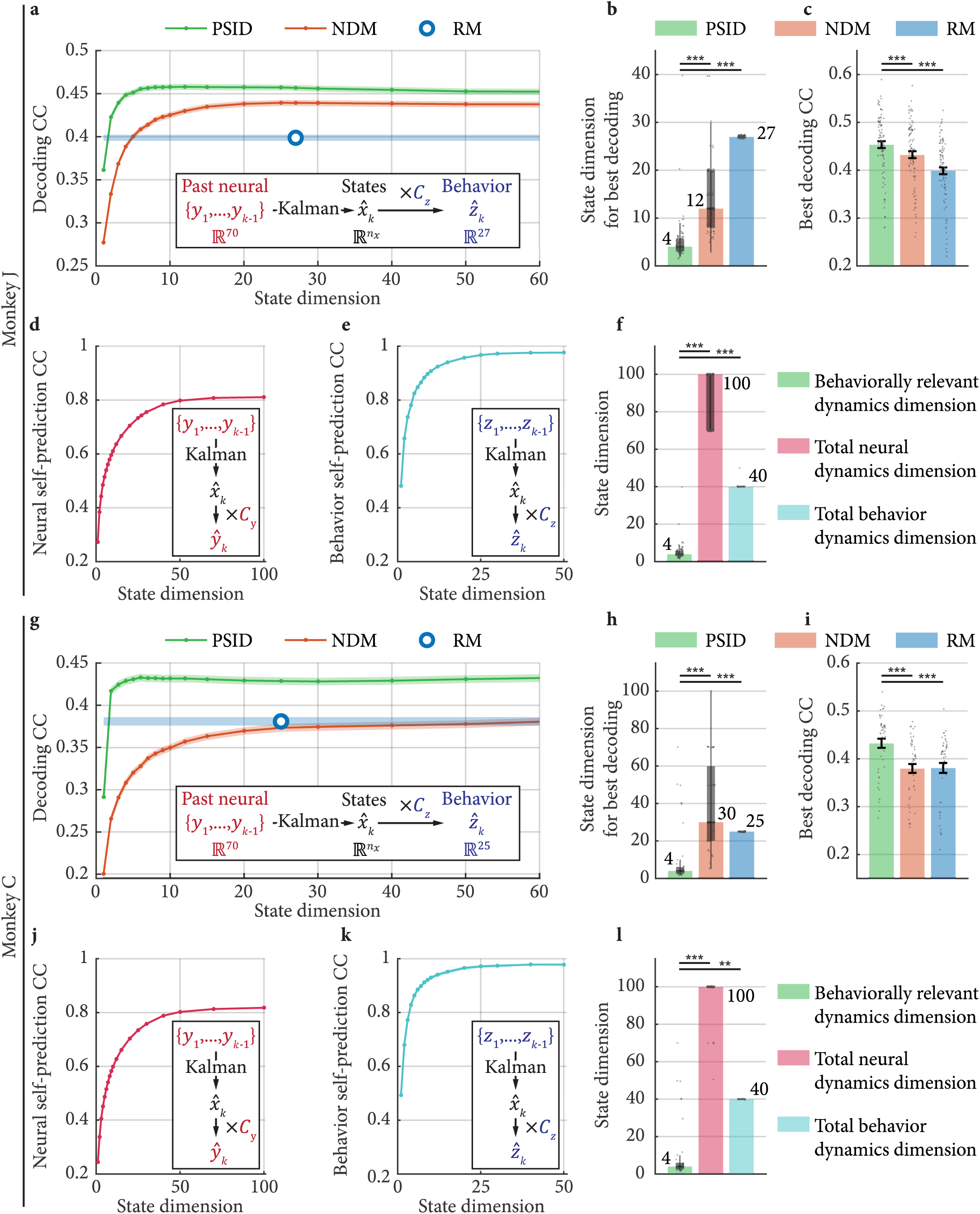
PSID reveals a markedly lower dimension for behaviorally relevant neural dynamics in the motor cortex during unconstrained naturalistic 3D reach, grasp and return movements. (**a**) Average joint angle decoding accuracy, i.e. cross-validated correlation coefficient (CC), as a function of the state dimension using PSID, NDM, and RM. Decoding CC is averaged across the datasets and the shaded area indicates the s.e.m. Dimensionality of neural activity (i.e. 70) and behavior (i.e. 27) are shown in a box along with the decoder structure. (**b**) The state dimension that achieves the best decoding in each dataset. Bars show the median (also written next to the bar), box edges show the 25^th^ and 75^th^ percentiles, and whiskers represent the minimum and maximum values (other than outliers). Outliers are the points that are more than 1.5 times the interquartile distance, i.e. the box height, away from the top and bottom of the box. All data points are shown. Asterisks indicate significance of statistical tests with *: P < 0.05, **: P < 0.005, ***: P < 0.0005, and n.s.: P > 0.05. (**c**) Best decoding CC in each dataset (state dimensions from (b)). For decoding, bars show the mean and whiskers show the s.e.m. (**d**) One-step-ahead self-prediction of neural activity (cross-validated CC), averaged across datasets. (**e**) Same as (d) for behavior. (**f**) The behaviorally relevant neural dynamics dimension (i.e. PSID result from (b)), total neural dynamics dimension (i.e. state dimension from (d)), and total behavior dynamics dimension (i.e. state dimension from (e)) for all datasets. (**g**)-(**l**) Same as (a)-(f), for monkey C.

### PSID requires fewer training samples

The previous results show that given the same training data, unlike NDM, PSID can identify the behaviorally relevant dynamics when used in the dimensionality reduction regime (i.e. with fewer latent states than the total number of latent states in the actual model, Fig. 2b for *n*_*x*_ < 16); and that even when the latent state dimension is as high as the actual model, PSID is more accurate than NDM in learning behaviorally relevant dynamics (Fig. 2b for *n*_*x*_ = 16). To further investigate how this PSID advantage depends on the training sample size, we evaluated each method when using different number of training samples. We found that RM and NDM in the dimensionality reduction regime could not learn behaviorally relevant dynamics even when training samples converged toward being unlimited (Supplementary Fig. 4a, b). Also importantly, even compared with NDM with a latent state dimension as high as the actual model, PSID achieved several orders of magnitude reduction in the number of training samples required to identify these dynamics because PSID prioritized them. In terms of both identifying behaviorally relevant eigenvalues and decoding behavior from neural activity, PSID required only about 0.2% of the training samples that NDM needed to achieve a similar accuracy (i.e. 500 times fewer; Supplementary Fig. 4). As training data in experiments is limited, this is another advantage of PSID, which aims to prevent the behaviorally relevant dynamics from being masked or confounded by the behaviorally irrelevant ones.

### PSID reveals a markedly lower dimensionality for behaviorally relevant neural dynamics in motor cortex

Given that PSID can prioritize learning of behaviorally relevant neural dynamics and dissociate them from behaviorally irrelevant ones, we used it to investigate the behaviorally relevant neural dynamics and their true dimensionality in large-scale motor cortical recordings during reach, grasp and return movements (Fig. 3, Methods). We found that PSID reveals the behaviorally relevant neural dynamics to be much lower-dimensional than would otherwise be concluded using standard methods (Fig. 3b, h), and that PSID identifies these dynamics more accurately than standard methods (Fig. 3a, c, g, i). To find the behaviorally relevant neural dynamics, we used PSID, NDM and RM to model neural features with various state dimensions (Fig. 3a, g). The dimension of behaviorally relevant neural dynamics is defined as the minimal state dimension required to best explain behavior using neural activity. To find this dimension from data, for each method and in each dataset, we found the smallest state dimension at which the best possible behavior decoding performance was achieved (Methods, Supplementary Fig. 5a, b). First, we found that the best possible decoding performance using PSID was significantly higher than the best possible decoding performance using both NDM and RM in both monkeys, suggesting that PSID more accurately learns behaviorally relevant neural dynamics (Fig. 3c, i; P < 10^−5^; one-sided signed-rank; *N*_*s*_ ≥ 48, Methods). Second, importantly, this best performance was achieved using a significantly smaller state dimension with PSID compared with NDM and RM—a median dimension of only 4 in both monkeys with PSID versus 12-30 with NDM and RM, or at least 3 times smaller (Fig. 3b, h; P < 10^−9^; one-sided signed-rank; *N*_*s*_ ≥ 48). Third, we confirmed with numerical simulations that PSID accurately estimates the true dimension of behaviorally relevant neural dynamics, whereas NDM overestimates it (Supplementary Fig. 5a, b). Finally, as a control analysis, we repeated NDM using the standard EM algorithm instead of the standard SID, and found similar results: PSID again achieved a significantly better decoding performance (P < 10^−9^; one-sided signed-rank; *N*_*s*_ ≥ 48) using significantly lower-dimensional latent states (P < 10^−7^; one-sided signed-rank; *N*_*s*_ ≥ 48). Together these results suggest that the behaviorally relevant motor cortical dynamics have a markedly lower dimension than is found by standard methods; PSID reveals this low dimension by more accurately learning behaviorally relevant neural dynamics and dissociating them from behaviorally irrelevant ones.

We next found that the dimensionality of the behaviorally relevant neural dynamics was much lower than that of neural dynamics or joint angle dynamics, suggesting that the low-dimensionality PSID finds is not simply because either neural or behavior dynamics are just as low-dimensional. To quantify the dimensionality of neural and behavior dynamics, we found the latent state dimension required to achieve the best self-prediction of neural or behavioral signals using their own past, and defined it as the total neural or behavior dynamics dimension, respectively (Methods). We confirmed in numerical simulations that this procedure correctly estimates the total latent state dimension in each signal (Supplementary Fig. 5c, d, e). First, for the neural features, we found that in both monkeys a median latent state dimension of at least 100 was required to achieve the best neural self-prediction (Fig. 3d, f, j, l), which is significantly larger than the behaviorally relevant neural dynamics dimension of 4 as revealed by PSID (P < 10^−18^; one-sided rank-sum; *N*_*s*_ ≥ 48). Second, for the behavior defined as joint angles, we found that in both monkeys a median latent state dimension of 40 was required to achieve the best behavior self-prediction (Fig. 3e, f, k, l), which is again significantly larger than the behaviorally relevant neural dynamics dimension of 4 as revealed by PSID (P < 0.004; one-sided rank-sum). Moreover, the *self-prediction* of behavior from its own past was much better than its decoding from neural activity (Fig. 3a, e, g, k) and reached an almost perfect CC of 0.98 for both monkeys (Fig. 3e, k), indicating that there are predictable dynamics in behavior that are not present in the recorded neural activity (corresponding to *ϵ*_*k*_ in Fig. 1). Taken together, these results suggest that beyond the low-dimensional behaviorally relevant neural dynamics extracted via PSID, both recorded neural activity and behavior have significant additional dynamics that are predictable from their own past but are unrelated to the other signal; PSID uniquely enables the dissociation of shared dynamics from the dynamics that are present in one signal but not the other (Fig. 1).

Finally, we found that the above results held irrespective of the exact behavioral signal. We repeated all the above analyses for the 3D position of hand and elbow taken as the behavioral signal (instead of joint angles) and found consistent results (Supplementary Fig. 6). PSID again revealed a significantly lower dimension for behaviorally relevant neural dynamics compared with NDM for both monkeys (P < 10^−6^; one-sided signed-rank; *N*_*s*_ ≥ 48) and achieved a significantly better decoding compared with NDM and RM (P < 10^−8^; one-sided signed-rank; *N*_*s*_ ≥ 48). Moreover, in both monkeys, the dimension of behaviorally relevant neural dynamics revealed by PSID was again significantly smaller than the dimension of dynamics in the recorded neural activity (P < 10^−18^; one-sided rank-sum) and in behavior (P < 0.004; one-sided rank-sum) as estimated based on their self-prediction.

### PSID reveals behaviorally relevant rotational dynamics that otherwise go unnoticed

Reducing the dimension of neural population activity and finding its low-dimensional representation are essential for visualizing and characterizing the relationship of neural dynamics to behavior^14,16,20–22^. We hypothesized that PSID would be particularly beneficial for doing this compared with standard NDM methods, because PSID can prioritize and directly dissociate the behaviorally relevant dynamics within neural activity. To test this hypothesis, we used PSID and NDM to extract a 2D representation of neural dynamics (Fig. 4), which is commonly done to visualize neural dynamics on planes^14,20–22^. We then compared the properties and the decoding accuracy of the extracted 2D dynamics. To do this, using both PSID and NDM, we fitted models with latent states of dimension 2 to neural activity during our naturalistic 3D reach, grasp and return task (Fig. 4a), estimated the latent states from neural activity using these models (Methods), and then plotted the two estimated latent states against each other during reach and return movement epochs (Fig. 4b, c, e, f).

**Figure 4.**
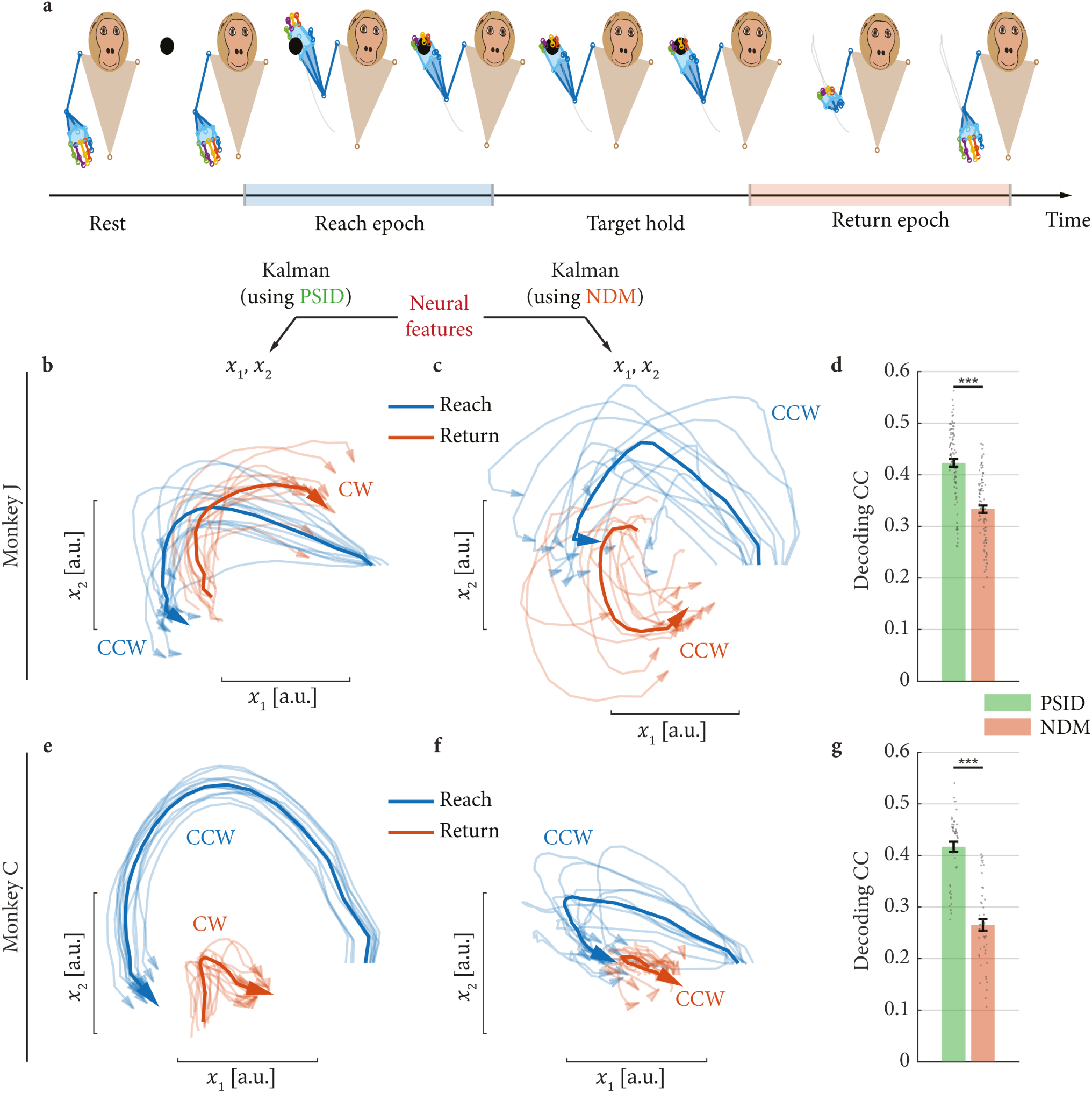
PSID reveals rotational neural dynamics with opposite direction during 3D reach and return movements, which is not found by standard methods. (**a**) Example reach and return epochs in the task defined as periods of movement toward the target and back from the target, respectively. Pictures are recreated using the 3D tracked markers and are from a view facing the monkey. (**b**) The latent neural state dynamics during 3D reach (blue) and return (red) movements found by PSID with 2D latent states (*n*_*x*_ = *n*_1_ = 2). We plot the states starting at the beginning of a reach/return movement epoch; the arrows mark the end of the movement epoch. Light lines show the average trace over trials in each dataset and dark lines show the overall average trace across datasets. The direction of rotation is noted by CW for clockwise or CCW for counter clockwise. States have arbitrary units (a.u.). (**c**) Same as (a) but using NDM with 2D latent states (*n*_*x*_ = 2). (**d**) Cross-validated correlation coefficient (CC) between the decoded and true joint angles, decoded with the latent states extracted using PSID and NDM in (a) and (b). Bars, whiskers and asterisks are defined as in Fig. 3c. (**e**)-(**g**) Same as (b)-(d), for monkey C.

We found that in both monkeys, both PSID and NDM extracted neural states that exhibited rotational dynamics. This suggests that our complex task with unconstrained naturalistic 3D reaches and grasps involves rotational motor cortical dynamics akin to what has been observed for reaching during other tasks, often involving 2D cursor control^14,20–22^. However, surprisingly, a clear difference emerged in the properties of rotations uncovered by PSID compared with NDM when we considered the dynamics during the return movement epochs. During the return epochs, the 2D neural dynamics extracted using PSID showed a rotation in the opposite direction of the rotation during the reach epochs (Fig. 4b, e). In contrast, similar to results from prior work^21^, neural dynamics extracted using NDM showed a rotation in the same direction during both reach and return epochs (Fig. 4c, f). As the behavior involves opposite directions of movement during reach and return epochs, these results intuitively suggest that PSID finds a low-dimensional mapping of neural population activity that is more behaviorally relevant (Fig 4a). To quantify this suggestion, we decoded the behavior using the low-dimensional latent states in each case. We found that the 2D latent states extracted using PSID explained the behavior significantly better than those extracted using NDM and led to significantly better decoding (Fig. 4d, g; P < 10^−9^; one-sided signed-rank; *N*_*s*_ ≥ 48). Moreover, the decoding accuracy using the PSID extracted 2D states was only 7% (Monkey J) or 4% (Monkey C) worse than the best possible PSID decoding whereas for NDM the decoding using 2D states was 23% (Monkey J) or 30% (Monkey C) worse than NDM’s best possible decoding (Fig. 3a, g). This indicates that while both types of rotational dynamics depicted in Fig. 4 exist in the high-dimensional manifold traversed by the neural activity, PSID extracted the 2D mapping that preserved the more behaviorally relevant neural dynamics (an illustrative example is provided in Supplementary Video 1). These results suggest that PSID can reveal low-dimensional behaviorally relevant neural dynamics that may otherwise be missed when using standard NDM methods.

Beyond the above 2D results, the marked advantage of PSID over NDM when performing dimensionality reduction held across all dimensions (Fig. 3a, g). At any given latent state dimension, PSID extracted a low-dimensional state that resulted in substantially better decoding compared with NDM (Fig. 3a, g). This suggests that even beyond a 2D dimensionality reduction for visualization, PSID could be used as a general dynamic dimensionality reduction method that preferentially preserves the most behaviorally relevant dynamics (Discussion).

Finally, as a control, we found that jPCA, which is another behavior agnostic method specifically designed for extracting rotational dynamics^20^, also extracted unidirectional rotations similar to NDM (Supplementary Fig. 7). As another control, we repeated NDM with standard EM algorithm instead of the standard SID and found that it again extracted very similar unidirectional rotations as those found with SID.

### PSID extracted dynamics are more informative of behavior for almost all joints

Previous results showed that on average across the arm and finger joints, PSID identified latent states that led to significantly better decoding of reach, grasp, and return behavior compared with states of the same (Fig. 3a, g) or even higher dimension obtained from NDM or from RM (Fig. 3c, i). We next found that this result held for almost all arm or finger joints separately as well and was not restricted to a limited set of joints (e.g. only finger joints). Computing the best decoding accuracy of each joint separately (Supplementary Fig. 8), we found that PSID achieved better decoding than NDM for all individual joints in both monkeys and that this difference was statistically significant in almost all joints (Supplementary Fig. 8b, d; P < 10^−4^ for all joints in monkey C and P < 10^−12^ for 25 of 27 joints in monkey J; one-sided signed-rank test; *N*_*s*_ = 240 and *N*_*s*_ = 455 for monkeys C and J, respectively). Moreover, PSID achieved significantly better decoding than RM for all 27 joints in monkey J (Supplementary Fig. 8b; P < 0.04 for each joint; one-sided signed-rank; *N*_*s*_ = 455) and for 24 of the 25 joints in monkey C (Supplementary Fig. 8d; P < 0.004 for each joint; one-sided signed-rank; *N*_*s*_ = 240), and similar decoding for 1 joint in monkey C (*P* = 0.27 two-sided signed-rank; *N*_*s*_ = 240). Additionally, the significantly better decoding in PSID was achieved using states of significantly lower dimension compared with NDM and RM (Supplementary Fig. 8a, c; P < 10^−90^; one-sided signed-rank; *N*_*s*_ ≥ 1200). Specifically, PSID used a median state dimension of 3 (monkey J) or 2 (monkey C) while NDM used a median state dimension of 8 (monkey J) or 15 (monkey C), and RM used a state dimension of 27 (monkey J) or 25 (Monkey C).

### PSID extracted dynamics are more informative of behavior for almost all recording channels across premotor, primary motor, and prefrontal areas

We found that PSID was extracting more behaviorally relevant information from each recording channel rather than performing an implicit channel selection by discarding some channels with no behaviorally relevant information. To distinguish between these alternatives, we repeated the modeling but this time using only the neural features from one channel at a time (Fig. 5). We found that for both monkeys, PSID achieved significantly better decoding of behavior in at least 96% and 98% of individual channels compared with NDM and RM, respectively (Fig. 5b, d; P < 0.05 for each channel; one-sided signed-rank; *N*_*s*_ ≥ 20). Moreover, PSID achieved this significant improvement in decoding while using significantly lower state dimensions than NDM and RM (Fig. 5a, c; P < 10^−68^; one-sided signed-rank; *N*_*s*_ ≥ 512). Specifically, PSID used a median state dimension of only 5 for both monkeys while NDM used a median state dimension of 15 (monkey J) or 20 (monkey C), and RM used a state dimension of 27 (monkey J) or 25 (Monkey C). Thus, while recording channels from different anatomical regions (including ipsilateral PMd and PMv coverage in monkey C) had different ranges of decoding accuracy (Fig. 5b, d), even channels with a relatively weak decoding saw an improvement in decoding accuracy when using PSID. These results suggest that almost all channels contained behaviorally relevant dynamics and PSID could more accurately model these dynamics leading to better decoding of behavior while also using lower-dimensional latent states.

**Figure 5.**
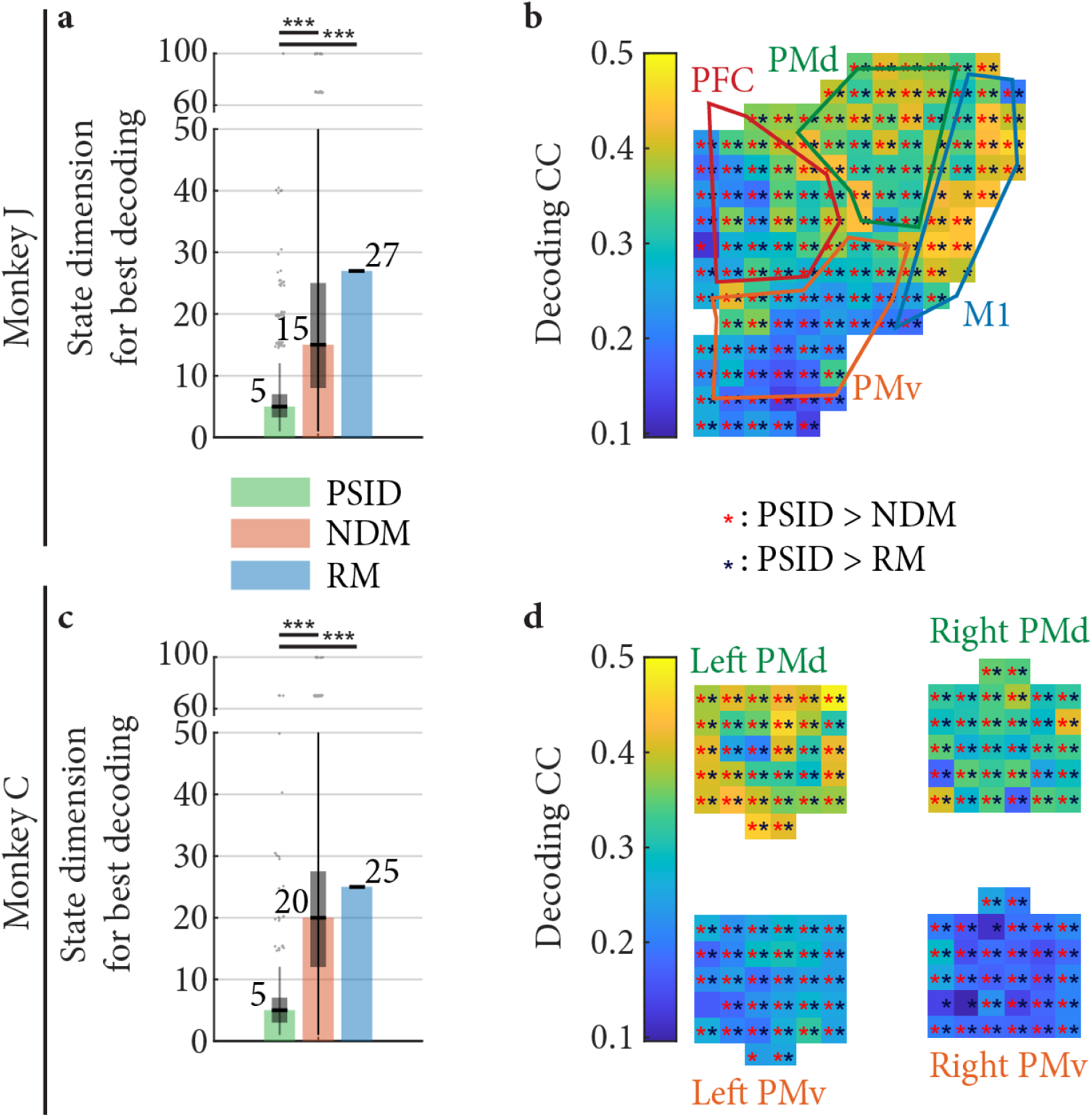
PSID more accurately identified the behaviorally relevant dynamics in each recording channel across premotor, primary motor, and prefrontal areas. (**a**)The state dimension used by each method to achieve the best decoding using the neural features from each recoding channel separately. For PSID and NDM, for each channel, the latent state dimension is chosen to be the smallest value for which the decoding CC reaches within 1 s.e.m. of the best decoding CC using that channel among all latent state dimensions. Bars, boxes and asterisks are defined as in Fig. 3b. (**b**) Cross-validated correlation coefficient (CC) between the decoded and true joint angles is shown for PSID. Asterisks mark channels for which PSID results in significantly (P < 0.05) better decoding compared with NDM (red asterisk) or RM (dark blue asterisk). The latent state dimension for each method is chosen as in (a). (**c**)-(**d**) Same as (a)-(b), for monkey C.

## Discussion

Here we develop and demonstrate a novel PSID algorithm for dissociating and modeling behaviorally relevant neural dynamics. Our simulations showed that compared with current methods, PSID learns the behaviorally relevant neural dynamics significantly more accurately, with markedly lower-dimensional latent states, and orders of magnitude fewer training samples. Our analyses on NHP motor cortical activity during an unconstrained 3D reach, grasp and return task confirmed these findings and revealed multiple new features of the underlying neural dynamics. First, PSID revealed the behaviorally relevant neural dynamics to be much lower-dimensional than implied by standard methods, and identified these dynamics more accurately as evident by better behavior decoding (Fig. 3). Second, PSID revealed distinct low-dimensional rotational dynamics in neural activity with opposite directions of rotation during reach and return epochs, which were more predictive of behavior than the alternative unidirectional rotational dynamics found by standard methods (Fig. 4). Finally, PSID resulted in significantly better decoding for almost any arm and finger joint angle (Supplementary Fig. 8) and for individual recording channels (Fig. 5). These results suggest that PSID can reveal low-dimensional behaviorally relevant neural dynamics that can otherwise go unnoticed.

The key idea in PSID was to ensure behaviorally relevant neural dynamics are not missed or confounded by prioritizing them in fitting the dynamic model. To do so, PSID models the neural activity as a latent SSM while prioritizing latent states that are informative of the behavior. Prior methods for NDM, including the standard SID or EM with linear dynamics^5,16,30,32^ as well as those with generalized linear dynamic systems (GLDS)^29,35^ or nonlinear dynamic models such as recurrent neural networks (RNN)^22^, are agnostic to behavior in fitting the dynamic model unlike PSID that takes behavior into account in fitting the dynamic model. Thus PSID can uncover important behaviorally relevant neural dynamics that may otherwise be discarded, such as the reversed rotational dynamics during return epochs in our task that were not revealed by NDM (Fig. 4, Supplementary Video 1).

Prior works have reported low-dimensional rotational neural dynamics during different tasks, often involving 2D control of a cursor^14,20–22^. Here we also found low-dimensional rotational dynamics during an unconstrained naturalistic 3D reach, grasp and return task—using PSID and NDM that have no supervision to try to do so as well as jPCA^20^ that aims to find rotations. However, while both NDM and PSID revealed rotations in neural dynamics during reach epochs, interestingly, the directions of the identified rotations were different in the return epochs between NDM and PSID. Similar to prior work applying NDM and jPCA to a center-out 2D cursor control task^21^, here NDM and jPCA extracted rotations in the same direction during reach and return epochs. In contrast, PSID extracted rotations that were in the opposite directions during reach and return epochs, and further were more behaviorally relevant (i.e. had significantly better behavior decoding accuracy, which was also close to the best decoding possible with even large latent state dimensions). This result demonstrates that while both the NDM- and PSID-extracted low-dimensional rotational dynamics existed in the high-dimensional neural activity (Supplementary Video 1), PSID revealed a low-dimensional mapping that preserved the behaviorally relevant components of neural dynamics. Future application of PSID to other behavioral tasks and brain regions may similarly reveal behaviorally relevant features of neural dynamics that may otherwise not be uncovered.

Our neural data was recorded from the motor cortical areas, which strongly encode movement related information and thus have long enabled motor brain machine interfaces^1,2,23^. Given this strong motor encoding, both RM, which models the dynamics of behavior agnostic to neural activity^2,23^, and NDM, which indiscriminately models all neural dynamics agnostic to behavior^21,29,30,32^, have been successful in decoding movement. Despite this strong encoding in motor cortical activity, PSID still resulted in significant improvements in decoding compared with standard methods and did so using smaller latent state dimensions (Fig. 3 and Supplementary Fig. 8). Our per channel analysis further showed that every channel contained behaviorally relevant information, which was better learned using PSID, thus resulting in decoding improvements (Fig. 5). Many brain functions such as memory^36^ and mood^5^ or brain dysfunctions such as epileptic seizures^7^ could have a more distributed or less targetable representation in neural activity. As a result, using PSID in such applications may prove even more beneficial since the activity is likely to contain more behaviorally irrelevant dynamics.

PSID can also be viewed as a dynamic dimensionally reduction method that provides a low-dimensional mapping of neural activity while preserving the behaviorally relevant information. PSID is a dynamic method since it models the temporal structure in neural activity (equation (1))—how it evolves over time. It can hence also aggregate information over time to optimally extract the latent brain state (Methods). Dynamic dimensionality reduction methods—i.e. methods that explicitly take into account temporal structure in extracting latent states such as Gaussian process factor analysis (GPFA)^35^ and SSM^5,16,21,29,30,32,34^—perform the dimensionality reduction only based on neural activity and are agnostic to behavior. In contrast, PSID enables taking behavior into account to ensure behaviorally relevant neural dynamics are accurately revealed. Thus, by focusing on behaviorally relevant neural dynamics, PSID can achieve a targeted dynamic dimensionality reduction that can be more suitable for studying neural mechanisms underlying a behavior of interest. For example, a multitude of prior works have reported that variables with 10-30 dimensions can sufficiently explain the information in motor cortical neural activity using dynamic (or non-dynamic) dimensionality reduction algorithms such as GPFA, RNN, and SSM^3,13,19,21,22,30,34,35^. However, unlike PSID, the algorithms used in these works did not aim to explicitly dissociate the behaviorally relevant parts of neural dynamics. Here, PSID revealed a markedly lower dimension for the behaviorally relevant neural dynamics of around 4, which was significantly lower that the dimension of 12-30 implied by the standard NDM approach (Fig. 3). This result demonstrates the utility of PSID in accurately estimating the dimensionality of behaviorally relevant neural dynamics, which is a fundamental sought-after question across domains of neuroscience^3,13,19^.

For datasets with discrete classes of behavioral conditions, several non-dynamic dimensionality reduction methods such as linear discriminant analysis (LDA)^16^ and demixed principal component analysis (dPCA)^25^ can take the discrete behavior classes into account and find a low dimensional projection of neural activity that is suitable for dissociating those classes^16^. However, unlike PSID, these methods are not applicable to continuous behavioral measurements such as movements. Further these methods cannot learn dynamic models and hence do not model the temporal patterns of neural activity or aggregate information over time, which is important especially in studying temporally structured behaviors such as unconstrained movements^2,23^ or speech^4^. Thus, PSID is a unique method that can enable dynamic dimensionality reduction by modeling temporal structure in neural population activity, apply to continuous valued behavioral measurements, and extract behaviorally relevant low-dimensional representations (i.e. latent states) for neural activity.

PSID uses a linear state-space model formulation in which both the latent state dynamics and the observation model are defined as linear functions of the latent state. A linear observation model is suitable for modeling continuous-valued observations such as the log-power features extracted from LFP signals in this work^5,29,32,37^. For spiking activity, some prior works have used a linear observation model with the spike counts in time windows of various lengths taken as the observation^2,21,23^, for which PSID is readily applicable. More recent studies have shown that using a GLDS framework with a nonlinear point process observation model for the binary spike events could provide a more accurate mathematical model in BMIs^38,39^. A variation of NDM using SID has been developed for GLDS models^34^ and an interesting area of future investigation is to generalize PSID to enable learning GLDS models with behaviorally relevant latent states from binary spike events. Moreover, given the growing interest in multi-scale modeling of simultaneous spike-field activity^29,37,40,41^, developing a multiscale version of PSID that can model observations from multiple modalities and timescales together would be another interesting area of future investigation.

In addition to serving as a new method to investigate the neural mechanisms of behavior, PSID may also help with future neurotechnologies for decoding and modulating behaviorally relevant brain states such as BMIs or closed-loop deep brain stimulation (DBS) systems^7^. While the motor representations in our datasets were strong, PSID could still help with decoding of behavior regardless of the latent state dimension. This decoding benefit may be even greater for brain states that are less strongly encoded or require recording neural activity from a more distributed brain network that is involved in various functions and thus exhibits more behaviorally irrelevant dynamics^3,9,12,42^. Further, PSID was able to identify a markedly lower-dimensional state that achieved close to maximal decoding accuracy. The identification of this low-dimensional behaviorally relevant state will be critical for developing model-based controllers^43^ to modulate various brain functions with electrical or optogenetic stimulation. This is because controllers designed for models with lower-dimensional states are generally more robust^44^. Finally, developing adaptive methods for latent state-space models that can track changes in behaviorally relevant dynamics, for example due to learning or stimulation-induced plasticity^2,45–48^, and can appropriately select the learning rate^49^ during adaptation are important future directions.

Here we described PSID as a tool for extracting and modeling behaviorally relevant dynamics from neural activity. In this application, neural activity is taken as the primary signal and behavior is taken as a secondary signal encoded by the primary signal. While this is the typical scenario of interest in neuroscience and neural engineering, the mathematical derivation of PSID does not depend on the nature of the two signals (Methods). For example, one could take behavior as the primary signal and neural activity as the secondary signal. If so, PSID would extract neural-activity-related dynamics from behavior and optionally also identify any additional behavioral dynamics not encoded in the recorded neural activity. Indeed, all numerical simulations reported in this work could be interpreted as having either neural activity or behavior as the primary signal and the other as the secondary signal. Beyond that, the two signals could even be generated by completely different sources. For example, in studying interpersonal neural and behavioral synchrony^50^ and social behavior^10^, applying PSID to neural and/or behavioral signals that are synchronously recorded from two individuals may enable extraction and modeling of common dynamics between the two. In general, when signals acquired from two systems are suspected to have shared dynamics (e.g. because they may be driven by common dynamic inputs), PSID can be used to extract and model the shared dynamics.

Taken together, the novel PSID modeling algorithm introduced in this work can serve as a tool to advance our understanding of how behaviorally observable brain functions are encoded in neural activity across broad tasks and brain regions. Also, PSID may prove to be particularly beneficial in studies of less strongly encoded brain functions involved in emotion, memory, and social behaviors.

## Methods

### Dynamic model

#### Model formulation

We used a linear state space dynamic model to describe the temporal evolution of neural activity and behavior as:

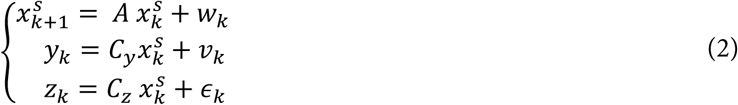

Here, *k* specifies the time index, 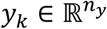 is the recorded neural activity, 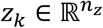 is the behavior (e.g., movement kinematics), 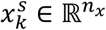 is the latent dynamic state variable that drives the recorded neural activity *y*_*k*_ and can also drive the behavior 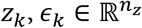 is a random process representing the dynamics in behavior that are not present in the recorded neural activity, and 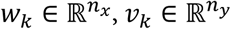 are zero-mean white noises that are independent of 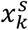, i.e. 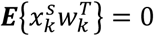 and 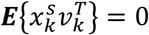 with the following cross-correlations:

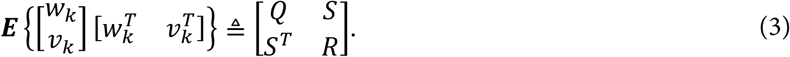

*ϵ*_*k*_ is a general random process denoting the variations of *z*_*k*_ that are not generated by 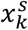 and thus are not present in the recorded neural activity. Thus, we only assume that *ϵ*_*k*_ is zero-mean and independent of 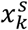, i.e. 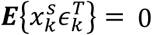 and the other noises, i.e. 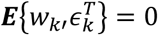 and 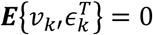 for any *k*′, but we do not make any assumptions about the dynamics of *ϵ*_*k*_. In fact, *ϵ*_*k*_ does not need to be white and can be any general non-white (colored) random process. Note that *ϵ*_*k*_ is also independent of *y*_*k*_ (since it is independent of 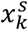 and *v*_*k*_), thus observing *y*_*k*_ does not provide any information about *ϵ*_*k*_. Due to the zero-mean assumption for noise statistics, it is easy to show that 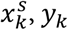, and *z*_*k*_ are also zero-mean, implying that in preprocessing, the mean of *y*_*k*_ and *z*_*k*_ should be subtracted from them and later added back to any model predictions if needed. The parameters *A, C*_*y*_, *C*_*z*_, *Q, R, S* fully specify the model in equation (2) (if statistical properties of *ϵ*_*k*_ are also of interest, another set of latent state-space parameters can be used to model it, Supplementary Note 1). There are other sets of parameters that can also equivalently and fully specify the model; Specifically, the set of parameters *A, C*_*y*_, *C*_*z*_, *G*_*y*_, ∑_*y*_, ∑_*x*_ with 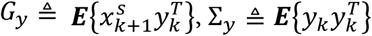, and 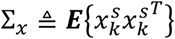 can also fully characterize the model and is more suitable for evaluating learning algorithms (Supplementary Note 2).

#### Definition of behaviorally relevant and behaviorally irrelevant latent states

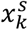 is a latent state that represents all dynamics in the neural activity *y*_*k*_, which could be due to various internal brain processes including the brain function of interest, other brain functions, or internal states. Without loss of generality, it can be shown (Supplementary Note 3) that equation (2) can be equivalently written in a different basis as

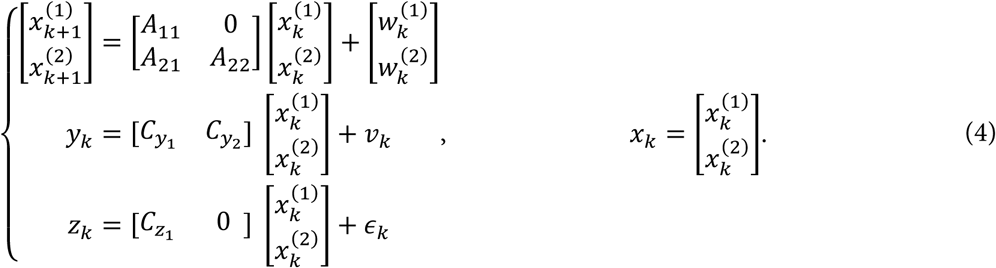

where 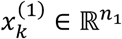 is the minimal set of states that affect behavior and whose dimension *n*_1_ is the rank of the behavior observability matrix (equation (42)). Thus, we refer to 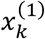 as the behaviorally relevant latent states and 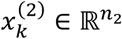 with *n*_2_ = *n*_*x*_ − *n*_1_ as the behaviorally irrelevant latent states. We interchangeably refer to the dimension of the latent states as the number of latent states (e.g. *n*_*x*_ is the total number of latent states or the total latent state dimension).

Equation (4) presents a general formulation of which special cases also include the models used in neural dynamics modeling (NDM) and representational modeling (RM). If we assume that all latent states can contribute to behavior (*n*_1_ = *n*_*x*_ and *n*_2_ = 0), equation (4) reduces to the linear SSM typically used to model the dynamics of neural activity in NDM^5,21,30,32,43^. If we further take *C*_*z*_ to be the identity matrix and *ϵ*_*k*_ = 0, the state will be set to the behavior *z*_*k*_ and equation (4) reduces to the linear SSMs used in RM^2,23^. Thus, if the assumptions of standard NDM (i.e. all latent states can drive both neural activity and behavior) or RM (i.e. behavior drives neural activity) hold better for a given dataset, PSID would still identify these standard models because the solution would still fall within the model in equation (4) used by PSID.

#### The learning problem

In the general learning problem, given training time series {*y*_*k*_ : 0 ≤ *k* < *N*} and {*z*_*k*_ : 0 ≤ *k* < *N*}, the aim is to find the dimension of the latent state *n*_*x*_ and all model parameters *A, C*_*y*_, *C*_*z*_, *G*_*y*_, ∑_*y*_, ∑_*x*_ that generate the data according to equation (2) or equivalently equation (4). Unlike prior work, here we critically require an identification algorithm that can dissociate the behaviorally relevant and irrelevant latent states, and can prioritize identification of the behaviorally relevant latent states (i.e. 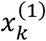 from equation (4)). Prioritizing behaviorally relevant latent states means that the algorithm would include the behaviorally relevant latent states in the model even when performing dimensionality reduction and thus identifying a model with fewer states than the true *n*_*x*_; this is typically the case given that training data is limited and neural dynamics are complex.

#### The decoding problem

Given the model parameters, the prediction (or decoding) problem is to provide the best estimate of *z*_*k*+1_ given the past neural activity {*y*_*n*_: 0 ≤ *n* ≤ *k*}. Given the linear state-space formulation of equation (2) and to achieve the minimum mean-square error, the best prediction of *y*_*k*+1_ using *y*_1_ to *y*_*k*_ and similarly the best prediction of *z*_*k*+1_ using *y*_1_ to *y*_*k*_—which we denote as *ŷ*_*k*+1|*k*_ and 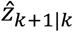, respectively—are obtained with the well-known recursive Kalman filter^51^ (Supplementary Note 4). By reformulating equation (2) to describe neural activity and behavior in terms of the latent states estimated by the Kalman filter, we can show that the best prediction of behavior using past neural activity is a linear function of the past neural activity (Supplementary Note 4). This key insight enables us to identify the model parameters via a direct estimation of the latent states through a projection of the future behavior onto the past neural activity (Supplementary Note 5).

### PSID: preferential subspace identification

We develop a novel learning algorithm, named preferential subspace identification (PSID), to identify the parameters of the dynamic model in equation (4) using training time series {*y*_*k*_ : 0 ≤ *k* < *N*} and {*z*_*k*_ : 0 ≤ *k* < *N*} while prioritizing the learning of the dynamics of *z*_*k*_ that are predictable from *y*_*k*_. The full algorithm is provided in Table 1. The detailed derivation is provided in Supplementary Note 5. In this section, we provide an overview of the derivation.

**Table 1.**
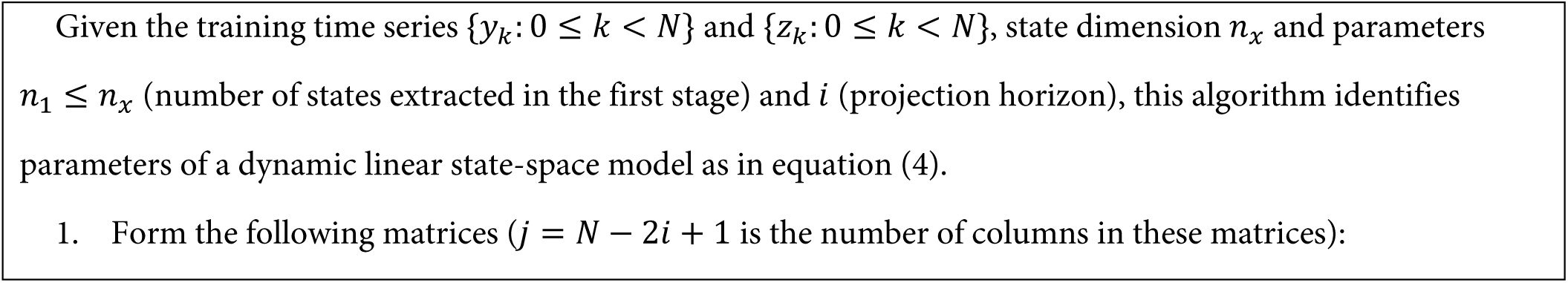
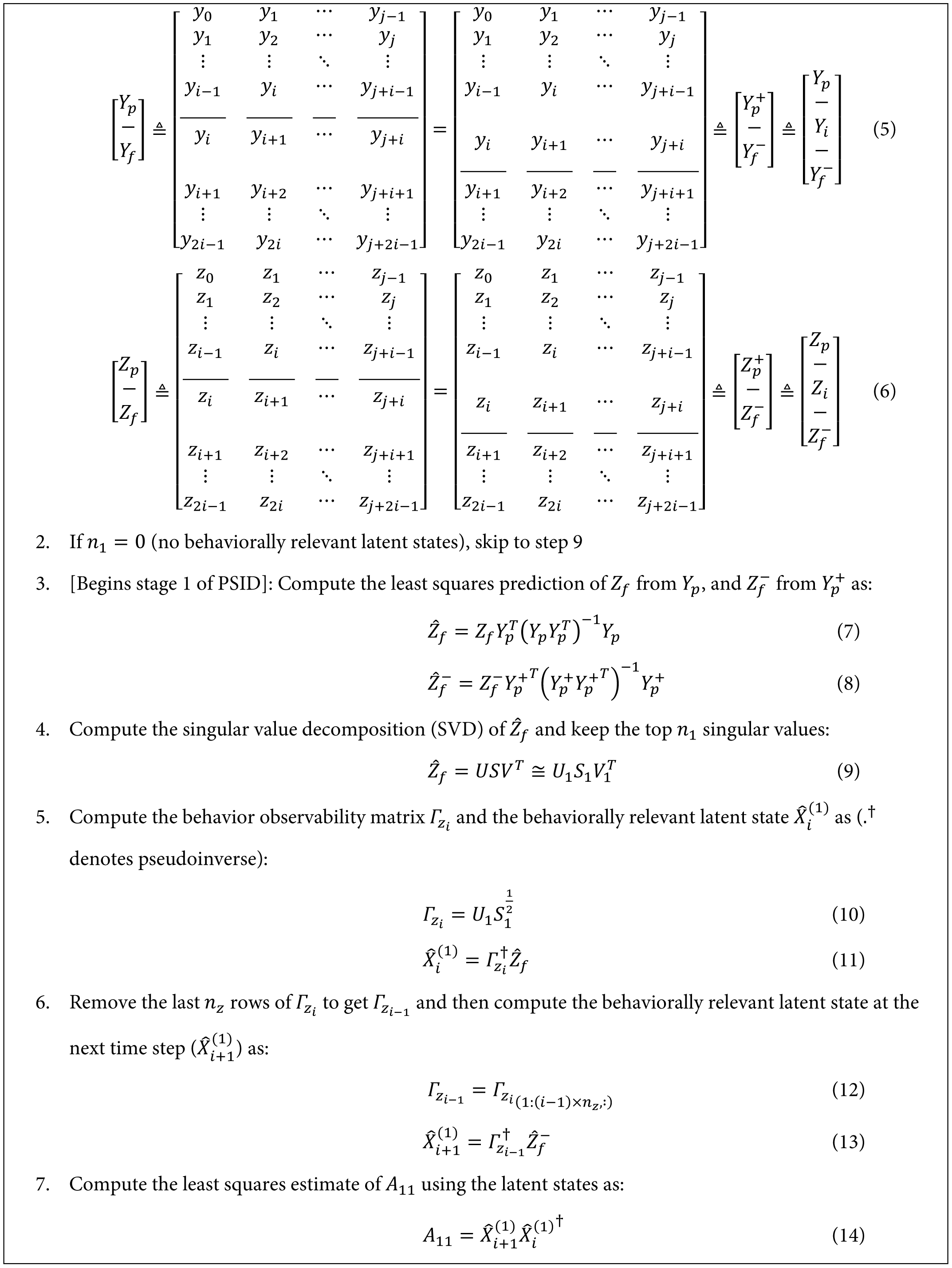
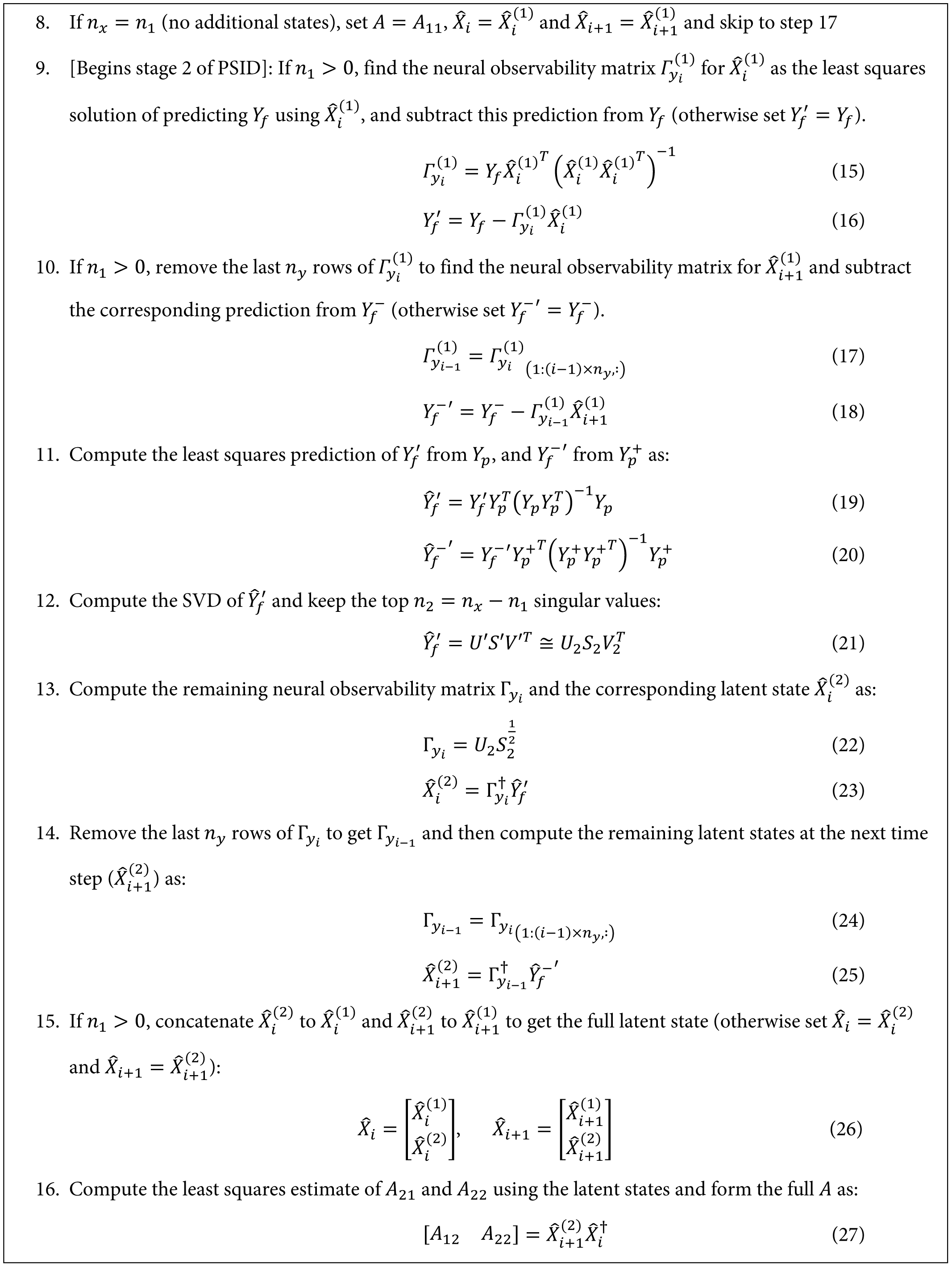
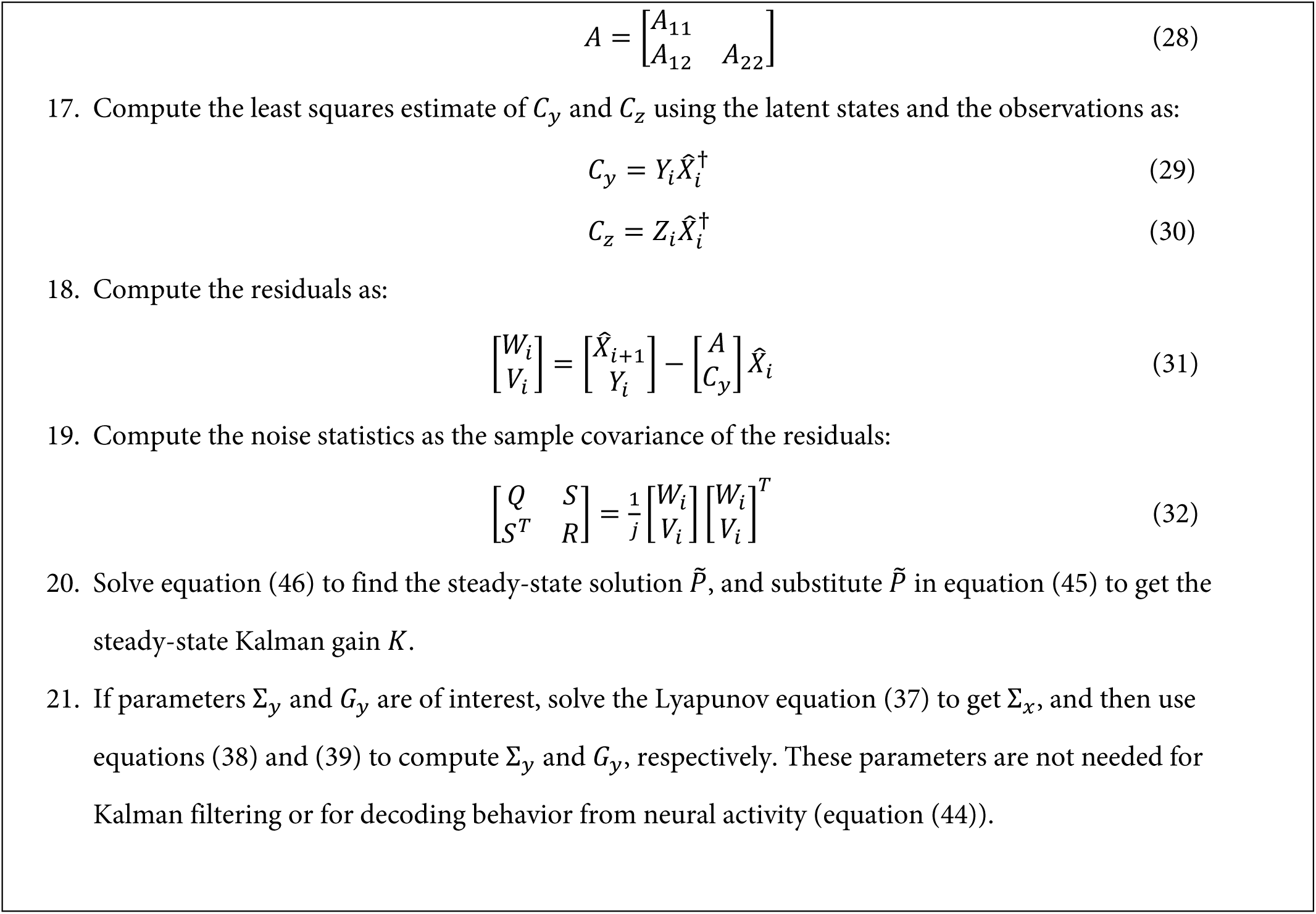
PSID: Preferential subspace identification algorithm.

PSID first extracts the latent states directly using the neural activity and behavior data, and then estimates the model parameters using the extracted latent states. The latent states are extracted in two stages: the first stage extracts behaviorally relevant latent states and the second stage, which is optional, extracts the remaining behaviorally irrelevant latent states. The first stage of PSID projects the future behavior (*Z*_*f*_) onto the past neural activity (*Y*_*p*_) (denoted as *Z*_*f*_/*Y*_*p*_ in Fig. 1b, equation (7)), which we can show extracts the behaviorally relevant latent states (Supplementary Note 5). The second stage of PSID first finds the part of the future neural activity that is not explained by the extracted behaviorally relevant latent states, i.e., does not lie in the subspace spanned by these states. This is found by subtracting the orthogonal projection of future neural activity onto the extracted behaviorally relevant latent states (equation (18)). This second stage then projects this unexplained future neural activity onto the past neural activity to extract the behaviorally irrelevant latent states (equation (19)). Overall, PSID provides a non-iterative closed-form solution for estimating the parameters of the model in equation (4) (Supplementary Note 5).

### Identification of model structure parameters for PSID and NDM

For both PSID and NDM, the total number of latent states *n*_*x*_ is a parameter of the model structure. When learning of all dynamics in the neural activity (regardless of their relevance to behavior) is of interest, we estimate the appropriate value for this parameter using the following cross-validation procedure. We fit models with different values of *n*_*x*_ and for each model, we compute the cross-validated accuracy of one-step-ahead prediction of neural activity *y*_*k*_ using its past (equation (44) in Supplementary Note 4). This is referred to as neural self-prediction to emphasize that the input is the past neural activity itself, which is used to predict the value of neural activity at the current time step. We use Pearson’s correlation coefficient (CC) to quantify the self-prediction (averaged across dimensions of neural activity). We then estimate the total neural latent state dimension *n*_*x*_ as the value that reaches within 1 s.e.m. of the best possible neural self-prediction accuracy among all considered latent state dimensions. As shown with numerical simulations, using this approach with PSID or standard SID^33,51^ for NDM accurately identifies the total number of latent states (Supplementary Fig. 3a-c and Supplementary Fig. 5c, e). We thus use this procedure to quantify the total neural dynamics dimension in NHP data (Fig. 3d, j). We also use the exact same procedure on the behavioral data using the behavior self-prediction to quantify the total behavior dynamics dimension in NHP data (Fig. 3e, k).

To learn a model with PSID with a given latent state dimension *n*_*x*_, we also need to specify another model structure parameter *n*_1_, i.e. the dimension of 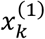 in equation (4). To determine a suitable value for *n*_1_, we perform an inner cross-validation within the training data and fit models with the given *n*_*x*_ and with different candidate values for *n*_1_. Among considered values for *n*_1_, we select the final value 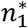 as the value of *n*_1_ that within the inner cross-validation in the training data, maximizes the accuracy for decoding behavior using neural activity (equation (44) in Supplementary Note 4). We quantify the decoding accuracy using CC (averaged across dimensions of behavior). As shown with numerical simulations, this approach accurately identifies *n*_1_ (Supplementary Fig. 3d, e). Thus, when fitting a model with any given latent state dimension *n*_*x*_ using PSID, unless otherwise noted, we determine *n*_1_ using an inner cross-validation as detailed above (Fig. 3a-c, Supplementary Fig. 5a, Supplementary Fig. 3a).

### Generating random models for numerical simulations

To validate the identification algorithms with numerical simulations, we generate random models with the following procedure. Dimension of *y*_*k*_ and *z*_*k*_ are selected randomly with uniform probability from the following ranges: 5 ≤ *n*_*y*_, *n*_*z*_ ≤ 10. The full latent state dimension is selected with uniform probability from 1 ≤ *n*_*x*_ ≤ 10 and then the number of states driving behavior (*n*_1_) is selected with uniform probability from 1 ≤ *n*_1_ ≤ *n*_*x*_. We then randomly generate matrices with consistent dimensions to be used as the model parameters *A, C*_*y*_, *C*_*z*_, *Q, R, S* (Supplementary Note 7). Specifically, the eigenvalues of *A* are selected randomly from the unit circle and *n*_1_ of them are then randomly selected to be used in the behaviorally relevant part of *A* (i.e. *A*_11_ in equation (4), Supplementary Note 7). Furthermore, noise statistics are randomly generated and then scaled with random values to provide a wide range of relative state and observation noise values (Supplementary Note 7). Finally, we generate a separate randomly generated SSM with a random number of latent states as the model for the independent residual behavior dynamics *ϵ*_*k*_ (Supplementary Note 7).

To generate a time-series realization with *N* data points from a given model, we first randomly generate an *N* data point white gaussian noise with the covariance given in equation (62) and assign these random numbers to *w*_*k*_ and *v*_*k*_. We then compute *x*_*k*_ and *y*_*k*_ by iterating through equation (2) with the initial value *x*_−1_ = 0. Finally, we generate a completely independent *N*-point time-series realization from the behavior residual dynamics model (see the previous paragraph) and add its generated behavior time series (i.e. *ϵ*_*k*_) to *C*_*z*_*x*_*k*_ to get the total *z*_*k*_ (equation (2)).

### Evaluation metrics for learning of model parameters in numerical simulations

A similarity transform is a revertible transformation of the basis in which states of the model are described and can be achieved by multiplying the states with any invertible matrix (Supplementary Note 2). For example, any permutation of the states is a similarity transform. Since any similarity transform on the model gives an equivalent model for the same neural activity and behavior (just changes the latent state basis in which we describe the model; Supplementary Note 2), we cannot directly compare the parameters of the identified model with the true model and need to consider all similarity transforms of the identified model as well. Thus, to evaluate the identification of model parameters, we first find a similarity transform that makes the basis of the latent states for the identified model as close as possible to the basis of the latent states for the true model. We then evaluate the difference between the identified and true values of each model parameter. Purely to find such a similarity transform, from the true model we generate a new realization with *q* = 1000*n*_*x*_ samples, which is taken to be sufficiently long for the model dynamics to be reflected in the states. We then use both the true and the identified models to estimate the latent state using the steady-state Kalman filter (equation (44)) associated with each model, namely 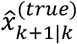 and 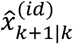. We then find the similarity transform that minimizes the mean-squared error between the two sets of Kalman estimated states as

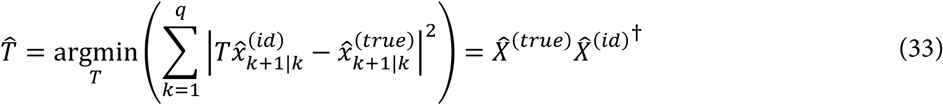

where 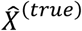 and 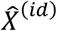 are matrices whose *k*th column is composed of 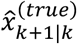 and 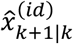, respectively. We then apply the similarity transform 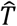 to the parameters of the identified model to get an equivalent model in the same basis as the true model. We emphasize again that the identified model and the model obtained from it using the above similarity transform are equivalent (Supplementary Note 2).

Given the true model and the transformed identified model, we quantify the identification error for each model parameter Ψ (e.g. *C*_*y*_) using the normalized matrix norm as:

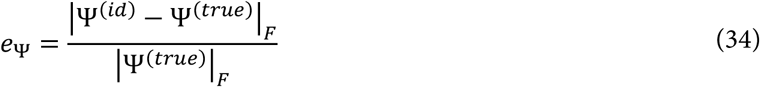

where |. |_*F*_ denotes the Frobenius norm of a matrix, which for any matrix Ψ = [*ψ*_*ij*_]_*n*×*M*_ is defined as:

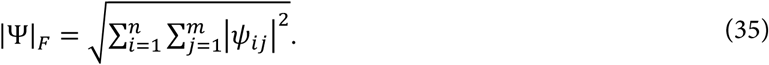

This concludes the evaluation of the identified model parameters.

### Evaluation metrics for learning of behaviorally relevant dynamics in numerical simulations

Both for numerical simulations and for NHP data, we use the cross-validated accuracy of decoding behavior using neural activity as a measure of how accurately the behaviorally relevant neural dynamics are learned. In numerical simulations, we also evaluate a more direct metric based on the eigenvalues of the state transition matrix *A*; this is because for a linear SSM, these eigenvalues specify the dynamical characteristics^52^. Specifically, we evaluate the identification accuracy for the eigenvalues associated with the behaviorally relevant latent states (i.e. eigenvalues of *A*_11_ in equation (4)). PSID identifies the model in the form of equation (4) and arranges the latent states such that the first block of *A* (i.e. *A*_11_ in equation (28)) is associated with the behaviorally relevant states (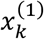 in equation (4)). Thus for PSID, we simply compute the eigenvalues of *A*_11_ and evaluate their identification accuracy. NDM identification methods do not specify which states are behaviorally relevant. So to find these states, we first apply a similarity transform to make the NDM identified *A* matrix block-diagonal with each complex conjugate pair of eigenvalues in a separate block (using MATLAB’s bdschur command followed by the cdf2rdf command). We then fit a linear regression from the states associated with each block to the behavior (using the training data) and sort the blocks by their prediction accuracy of behavior *z*_*k*_. The behaviorally relevant eigenvalues are then taken to be the top *n*_1_ eigenvalues that result in the most accurate prediction of *z*_*k*_.

Finally, given the true behaviorally relevant eigenvalues and the identified behaviorally relevant eigenvalues, we find the closest pairing of the two sets (by comparing all possible pairings), put the true and the associated closest identified eigenvalues in two vectors, and compute the normalized eigen value detection error using equation (34).

When evaluating the identified eigenvalues for models with a latent state dimension that is smaller than the true *n*_1_ (for example in Fig. 2), we add zeros instead of the missing eigenvalues since a model with fewer latent states is equivalent to a model with more latent states that are always equal to zero and have eigenvalues of zero associated with them.

### Identification of the dimensionality for behaviorally relevant neural dynamics

To estimate the dimensionality of the behaviorally relevant neural dynamics, we seek to find the minimal number (i.e., dimension) of latent states that is sufficient to best describe behavior using neural activity. To do this, for each method, we fit models with different values of state dimension *n*_*x*_, and compute the cross-validated accuracy of decoding behavior using neural activity (equation (44) in Supplementary Note 4). We use Pearson’s correlation coefficient (CC), averaged across behavior dimensions, to quantify the decoding accuracy. We then estimate the dimension of the behaviorally relevant neural dynamics as the smallest latent state dimension that reaches within 1 s.e.m. of the best possible cross-validated decoding accuracy among all considered latent stateas the smallest latent state dimension that reaches within 1 s.e.m. of the best possible cross-validated behavior decoding accuracy as described above (Fig. 3a-c).

### Recordings and task setup in non-human primates

Neural activity was recorded in two adult Rhesus macaques while the subjects were performing naturalistic reach, grasp, and return movements in a 3D space^37,53^. All surgical and experimental procedures were performed in compliance with the National Institute of Health Guide for Care and Use of Laboratory Animals and were approved by the New York University Institutional Animal Care and Use Committee. In Monkey J, neural activity was recorded from 137 electrodes on a micro-drive (Gray Matter Research, USA) covering parts of primary motor cortex (M1), dorsal premotor cortex (PMd), ventral premotor cortex (PMv), and prefrontal cortex (PFC) on the left hemisphere and in monkey C, activity was recorded from 128 electrodes on four thirty-two electrode microdrives (Gray Matter Research, USA) covering PMd and PMv on both left and right hemispheres. Using 3D tracked reflective markers, the movement of various points on the torso, chest, right arm, hand and fingers were tracked. These markers were used to extract the angle of the 27 (monkey J) or 25 (monkey C) joints of the upper-extremity, consisting of 7 joints in the shoulder, elbow, wrist, and 20 (monkey J) or 18 (monkey C) joints in fingers (4 in each, except 2 missing finger joints in monkey C)^53,54^. We analyzed the neural activity during 7 (monkey J) or 4 (monkey C) recording sessions. For most of our analyses (unless otherwise specified), to further increase the sample size, we randomly divided the electrodes into non-overlapping groups of 10 electrodes and performed modeling in each group separately. We refer to each random electrode group in each recording session as one dataset.

To model the recorded local field potentials (LFP), we performed common average referencing (CAR) and then as the neural features, extracted signal log-powers (i.e. in dB units) from 7 frequency bands^37,55^ (theta: 4-8 Hz, alpha: 8-12 Hz, low beta: 12-24 Hz, high beta: 24-34 Hz, low gamma: 34-55 Hz, high-gamma 1: 65-95 Hz, and high gamma 2: 130-170 Hz) within sliding 300ms windows at a time step of 50ms using Welch’s method (using 8 sub windows with 50% overlap)^56^. The extracted features were taken as the neural activity time series *y*_*k*_ (*y*_*k*_ ∈ ℝ^70^ in each dataset). Unless otherwise noted, the behavior time series *z*_*k*_ was taken as the joint angles at the end of each window (*z*_*k*_ ∈ ℝ^27^ in monkey J and *z*_*k*_ ∈ ℝ^25^ in monkey C).

### Cross-validated model evaluation and statistical tests on NHP neural datasets

For each method, we performed the model identification and decoding within a 5-fold cross-validation and as the performance metric for predicting behavior, we computed the cross-validated correlation coefficient between the true and predicted joint angles. For all methods, in each cross-validation fold, we first z-scored each dimension of neural activity and behavior based on the training data to ensure that learning methods do not discount any behavior or neural dimensions due to a potentially smaller natural variance. In fitting the models with PSID, for each latent dimension *n*_*x*_, unless specified otherwise, *n*_1_ was selected using a 4-fold inner cross-validation within the training data. For PSID and standard SID^33,51^, a horizon parameter of *i* = 5 was used in all analyses, except for per channel analyses (Fig. 5) where a horizon of *i* = 20 was used due to the smaller neural feature dimension. For the control analyses with NDM, we used the EM algorithm^57,58^.

We used the Wilcoxon signed-rank or rank-sum for all paired and non-paired statistical tests, respectively. To correct for multiple-comparisons when comparing the performance of methods for different joints or channels, we corrected the P-values within the test data using the False Discovery Rate (FDR) control^59^.

## Data availability

The data used to support the results are available upon reasonable request from the corresponding author.

## Code availability

The code for the PSID algorithm is available from the corresponding author and will be available online at https://github.com/ShanechiLab/PSID.

## Author contributions

O.G.S. and M.M.S. conceived and developed the new PSID algorithm. O.G.S. performed all the analyses. B.P. provided all the non-human primate data. O.G.S. and M.M.S. wrote the manuscript with input from B.P.

## Supplementary Figures

**Supplementary Figure 1.**
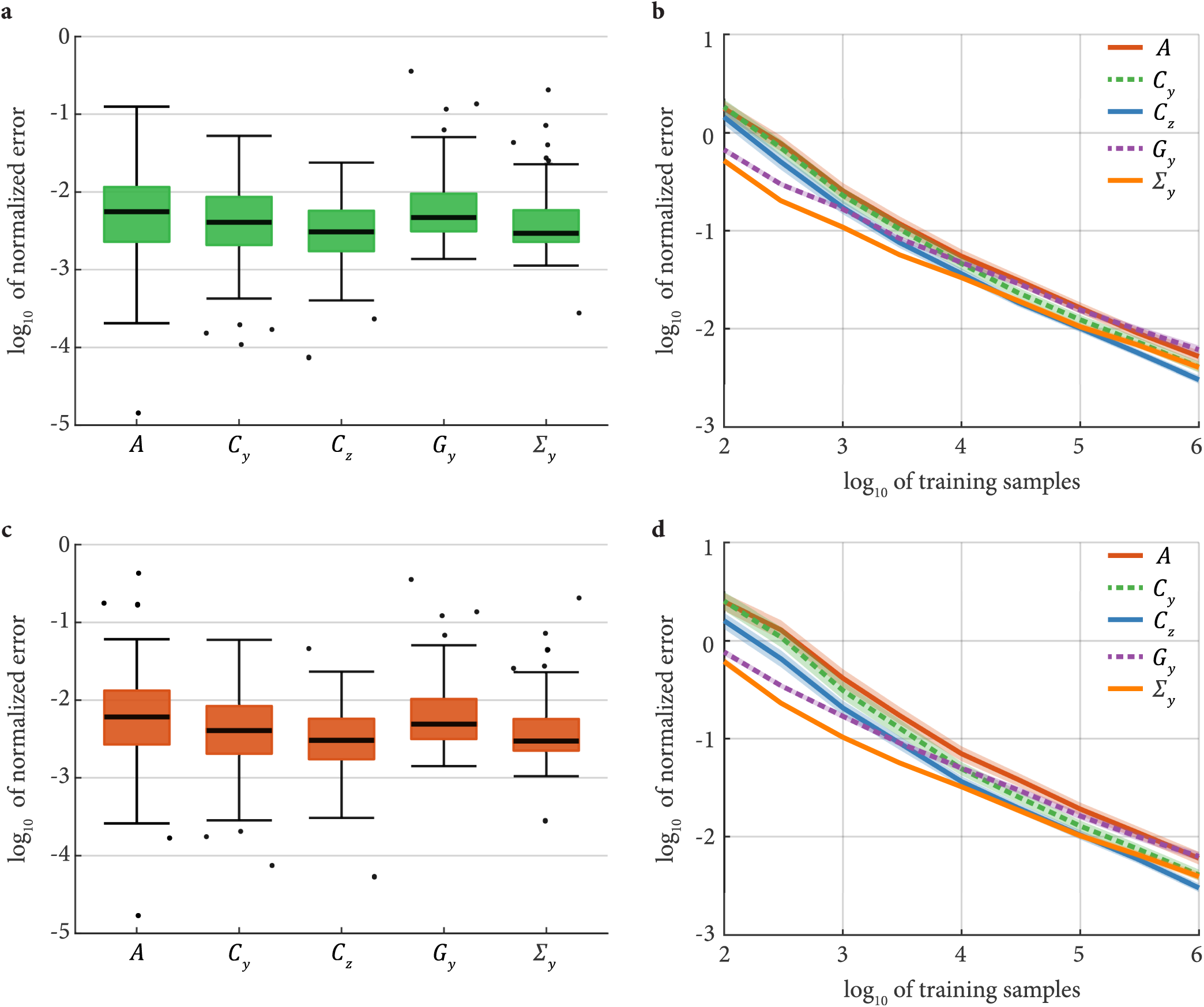
PSID correctly learns model parameters at a rate of convergence similar to that of SID while also being able to prioritize behaviorally relevant dynamics. **(a)** Normalized error for identification of each model parameter using PSID (with 10^6^ training samples) across 100 random simulated models. Each model had randomly selected state, neural activity, and behavior dimensions as well as randomly generated parameters (Methods). The parameters *A, C*_*y*_, *C*_*z*_ from equation (2) together with the covariance of the neural activity 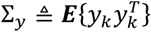 and the cross-covariance of the neural activity with the latent state 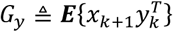 fully characterize the model (Methods). The horizontal dark line on the box shows the median, box edges show the 25th and 75th percentiles, whiskers represent the minimum and maximum values (other than outliers) and the dots show the outlier values. Outliers are defined as in Fig. 3. Using 10^6^ samples, all parameters are identified with a median error smaller than 1%. **(b)** Normalized error for all parameters as a function of the number of training samples for PSID. The normalized error consistently decreases as more samples are used for identification. Solid line shows the average log_10_ of the normalized error and the shaded area shows the s.e.m. **(c)-(d)** Same as (a)-(b), shown for the standard SID algorithm. The rate of convergence for both PSID and SID, and for all parameters is around 10 times smaller error for 100 times more training samples (i.e. slope of −0.5 on (b), (d)).

**Supplementary Figure 2.**
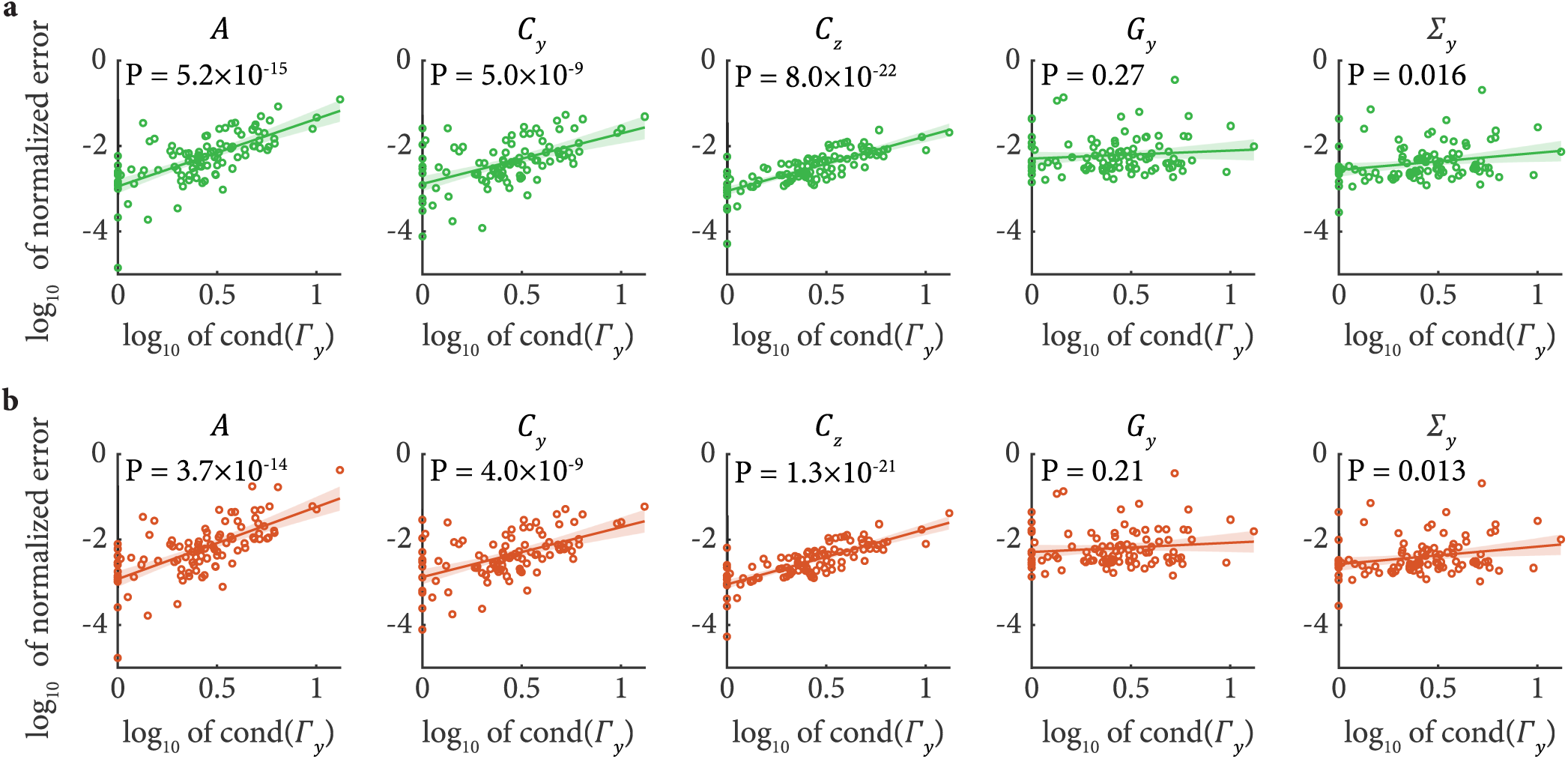
Identification error is larger for models that are closer to unobservability and thus inherently harder to identify. **(a)** Normalized error for each parameter (identified with PSID using 10^6^ training samples) for the 100 random simulated models in Supplementary Fig. 1 is shown as a function of the condition number of the neural observability matrix Γ_*y*_ for the model, which is defined as the ratio of its largest to its smallest singular values (Methods). The P-value for Pearson’s correlation coefficient between log_10_ cond(Γ_*y*_) and log_10_ of normalized error is shown on each plot (number of data points is 100). The green line shows the least squares solution to fitting a line to the data points and the shaded area shows the associated 95% confidence interval. The condition number of the neural observability matrix for each model is significantly correlated with the identification error for the three model parameters (i.e. *A, C*_*y*_, and *C*_*z*_) that have the widest range of identification errors (as seen from Supplementary Fig. 1a). As a model gets closer to being unobservable and more difficult to identify, the condition number for the observability matrix increases. Thus this result indicates that the models for which these three parameters were poorly estimated were closer to being unobservable and thus were inherently more difficult to identify given the same number of training samples. **(b)** Same as (a) for SID, which similarly shows relatively larger error for models that are inherently less observable.

**Supplementary Figure 3.**
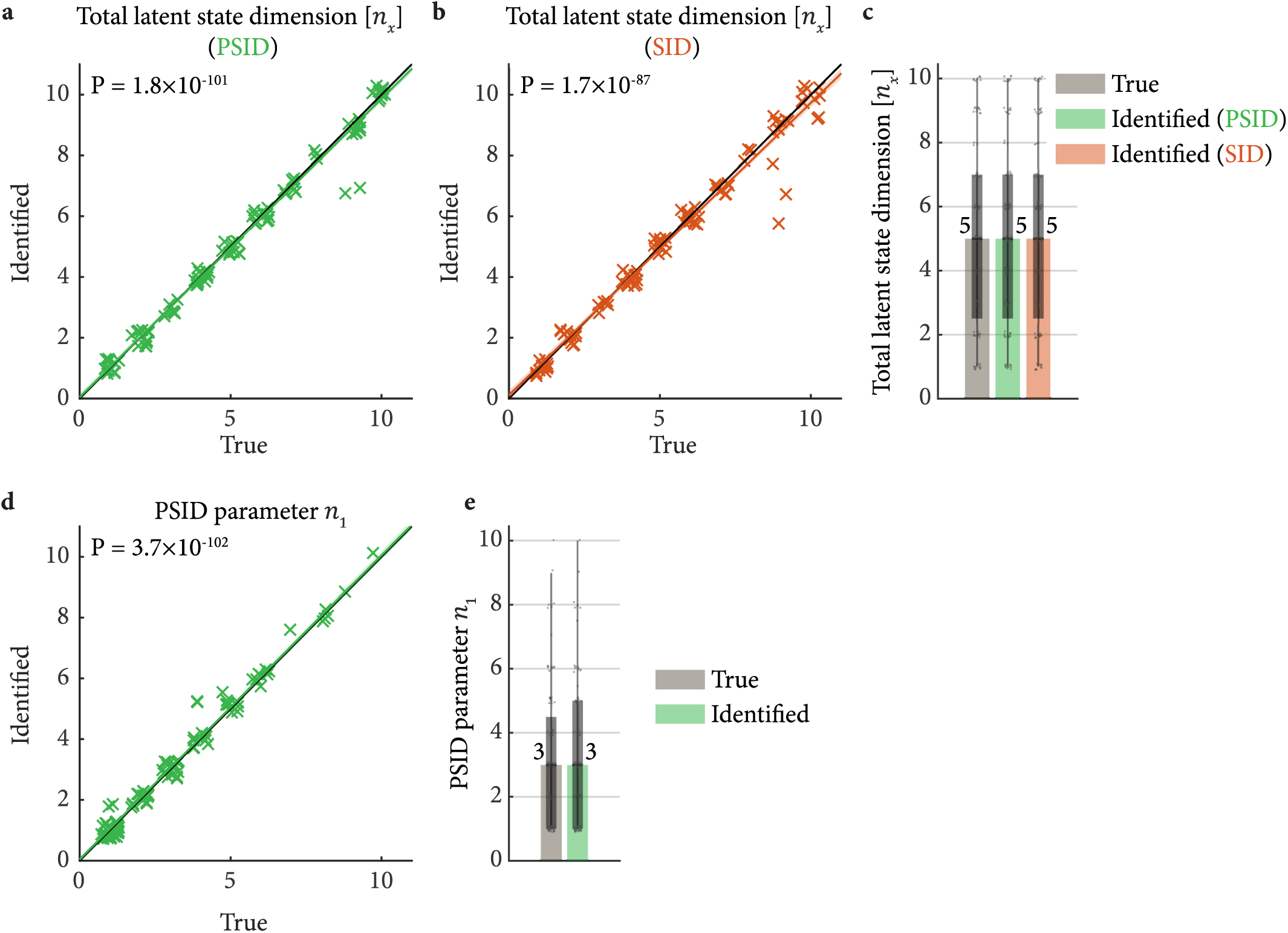
Model structure parameters can be accurately estimated using cross-validation. **(a)** Detection of the total latent state dimension (*n*_*x*_) using cross-validation is shown for numerical simulations. We estimate *n*_*x*_ by considering candidate values of *n*_*x*_ and selecting the value whose associated model reaches (within 1 s.e.m. of) the best neural *self-prediction* (predicting *y*_*k*_ using its past values) among all candidate values (Methods). The Pearson’s correlation P-value between the true and identified values is shown on the plot. The colored line and shaded area are defined as in Supplementary Fig. 2. **(b)** Same as (a), for detection of *n*_*x*_ using cross-validation in standard SID. **(c)** The distribution of true and identified values of *n*_*x*_ from (a)-(b) is shown as a box plot. Bars and boxes are defined as in Fig. 3b. All data points are shown. **(d)** Same as (a), for detection of the PSID parameter *n*_1_ (Methods). (e) The distribution of true and identified values of *n*_1_ from (d) shown as a box plot. The true and identified *n*_*x*_ and *n*_1_ are always integer values, so for better visualization and to avoid having multiple points at the exact same location on the plots a random small displacement has been added to each point.

**Supplementary Figure 4.**
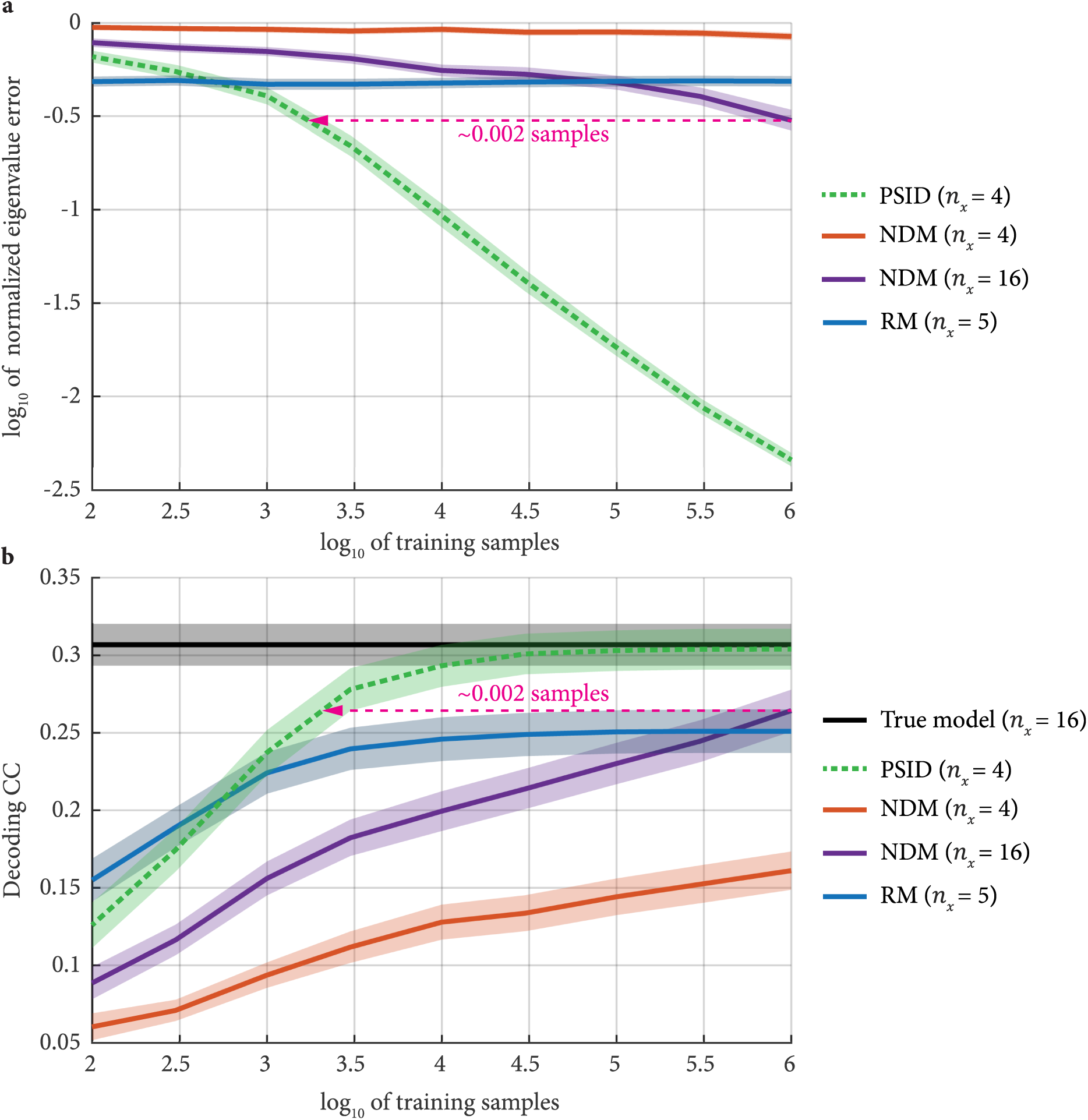
RM and NDM with the same latent state dimension as PSID cannot achieve a comparable performance to PSID even with unlimited training samples, and PSID requires orders of magnitude fewer samples to achieve the same performance as an NDM with a larger latent state dimension. **(a)** Normalized eigenvalue error is shown for 100 random simulated models when using RM, PSID, or NDM with similar or larger latent state dimension. Solid lines show the average across the 100 models, and the shaded areas show the s.e.m. For RM, the state dimension is the behavior dimension (here *n*_*z*_ = 5). **(b)** Cross-validated behavior decoding CC for the models in (a). Optimal decoding using the true model is shown as black. For NDM with 4 latent states (i.e. in the dimension reduction regime) and RM, eigenvalue identification and decoding accuracies plateaued at some final value below that of the true model and stopped improving with further addition of training samples, indicating that the asymptotic performance of having unlimited training samples has been reached. Even for an NDM with a latent state dimension as large as the true model (i.e. not performing any dimension reduction and using *n*_*x*_ = 16), (i) NDM was inferior in performance compared with PSID with a latent state dimension of only 4 when using the same number of training samples, and (ii) NDM required orders of magnitude more training samples to reach the performance of PSID with the smaller latent state dimension. Parameters are randomized as in Methods except the state noise (*w*_*t*_), which is 100 times smaller (i.e. −3 ≤ *α*_1_ ≤ −1), and the behavior signal-to-noise ratio, which is 10 times smaller (i.e. −1 ≤ *α*_3_ ≤ +1), both adjusted to make the decoding performances more similar to the NHP results.

**Supplementary Figure 5.**
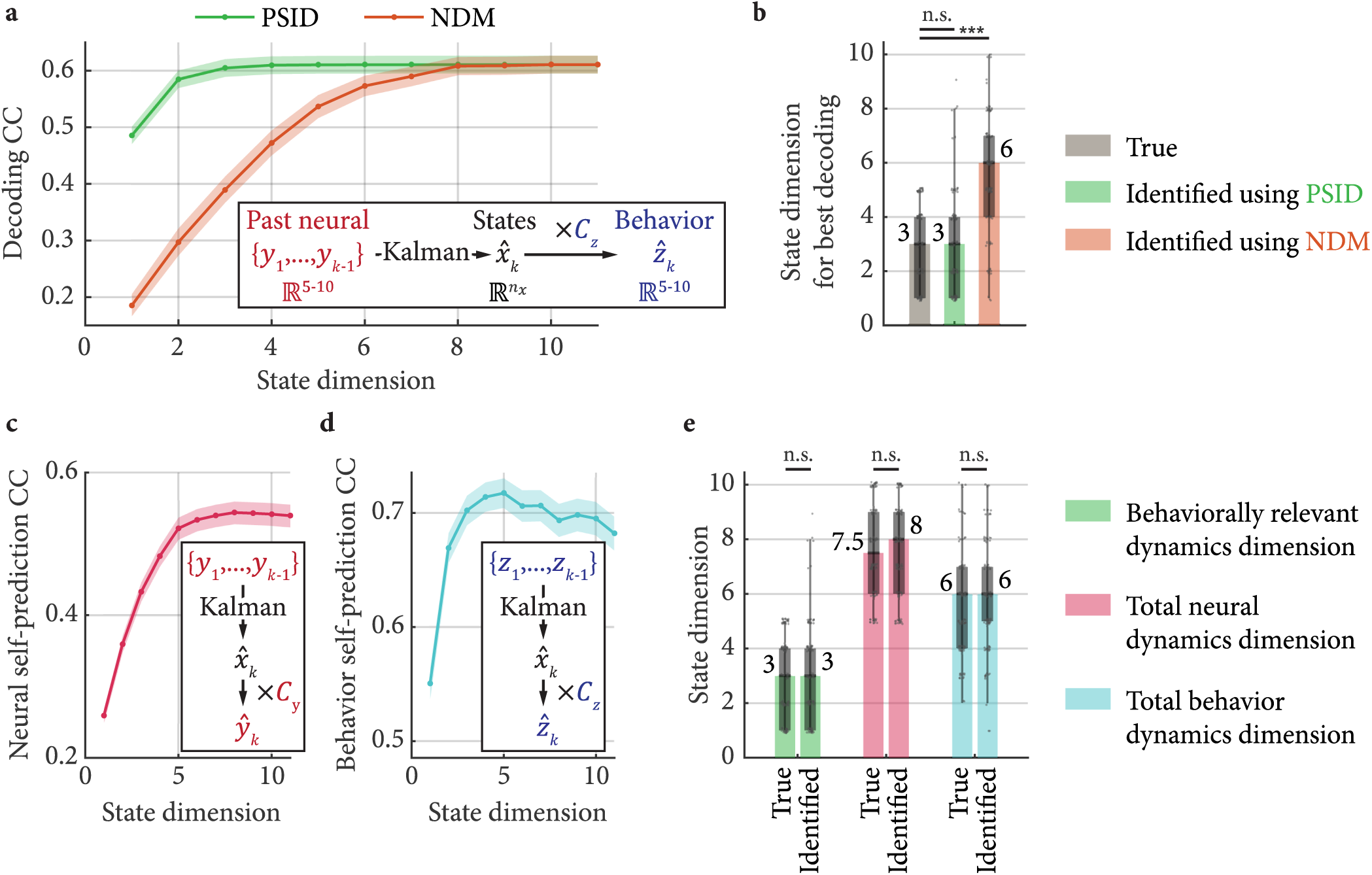
PSID can accurately estimate the behaviorally relevant dynamics dimension, as well as the total neural dynamics dimension and the total behavior dynamics dimension in simulations. **(a)** Cross-validated behavior decoding correlation coefficient (CC) as a function of latent state dimension using PSID and NDM within numerical simulations. Decoding CC is averaged across 100 random simulated models and the shaded area indicates the s.e.m. In each model, a random number of neural states were behaviorally irrelevant (Methods). **(b)** The behaviorally relevant neural dynamics dimension identified using PSID and NDM. This number is identified for each model as the smallest state dimension for which the CC reaches the best decoding performance. Bars, boxes and asterisks are defined as in Fig. 3b. While PSID accurately identifies the behaviorally relevant dynamics dimension, NDM overestimates it. **(c)** One-step-ahead self-prediction of neural activity (cross-validated CC) as a function of latent state dimension. To compute the self-prediction, SID (i.e., PSID with *n*_1_ = 0) is always used for modeling since dissociation of behaviorally relevant states is not needed. **(d)** Same as (c) for one-step-ahead self-prediction of behavior. **(e)** True and identified values for behaviorally relevant neural dynamics dimension (PSID results from (b)), the total neural dynamics dimension (identified as the state dimension for best neural self-prediction from (c)) and the total behavior dynamics dimension (identified as the state dimension for best behavior self-prediction from (d)). These results confirm with numerical simulations that our approach for identifying the total neural and behavior dynamics dimensions correctly estimates these numbers, and that PSID accurately identifies the behaviorally relevant dynamics dimension from data. Consequently, the same cross-validation approach is used in Fig. 3 for the real NHP data to compute the dimensions.

**Supplementary Figure 6.**
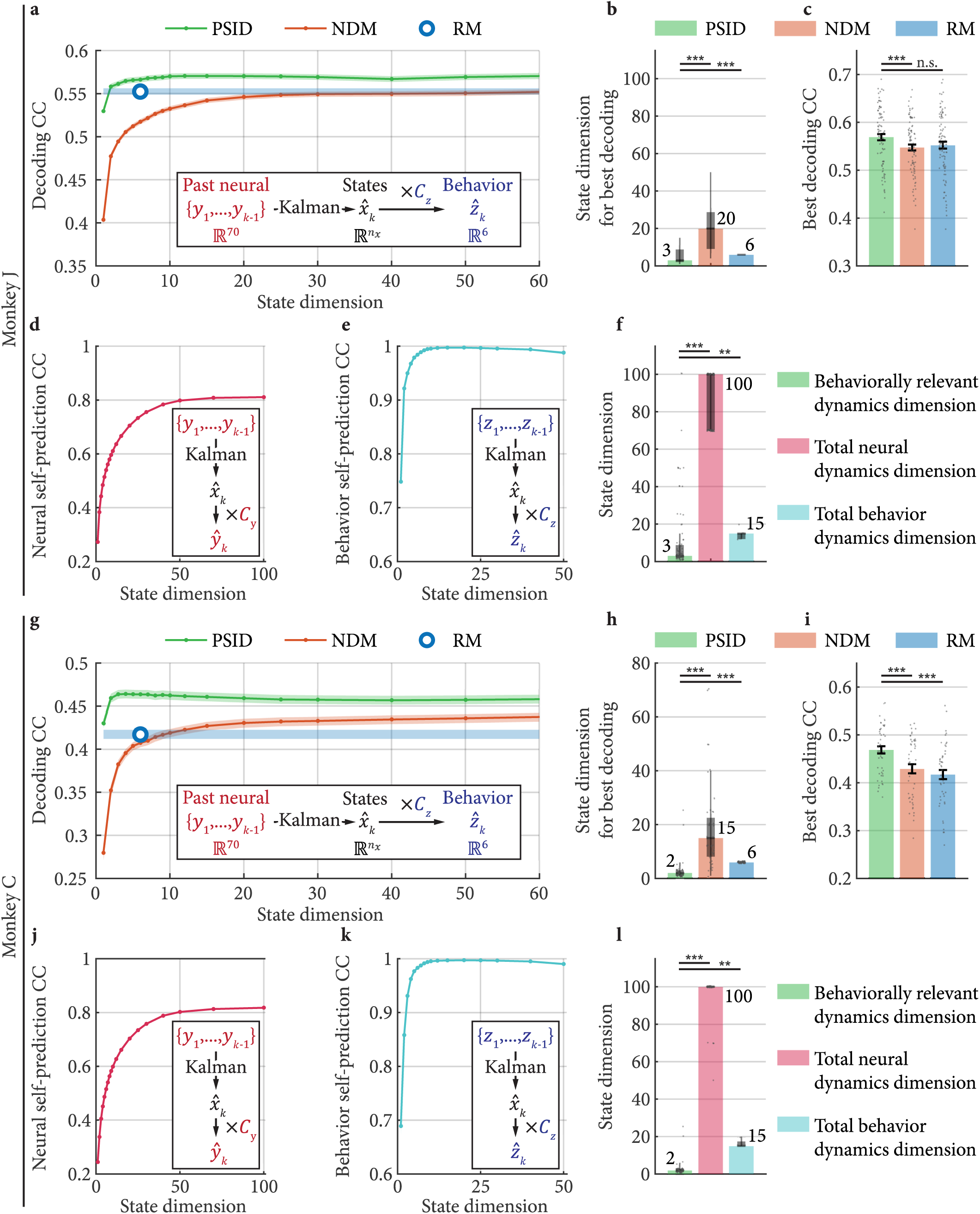
PSID again reveals a markedly lower dimension for behaviorally relevant neural dynamics in the motor cortex when behavior is taken as the 3D end-point position (of hand and elbow) instead of the joint angles. Notation is the same as in Fig. 3, but this time for behavior taken as the 3D position of hand and elbow (*n*_*z*_ = 6).

**Supplementary Figure 7.**
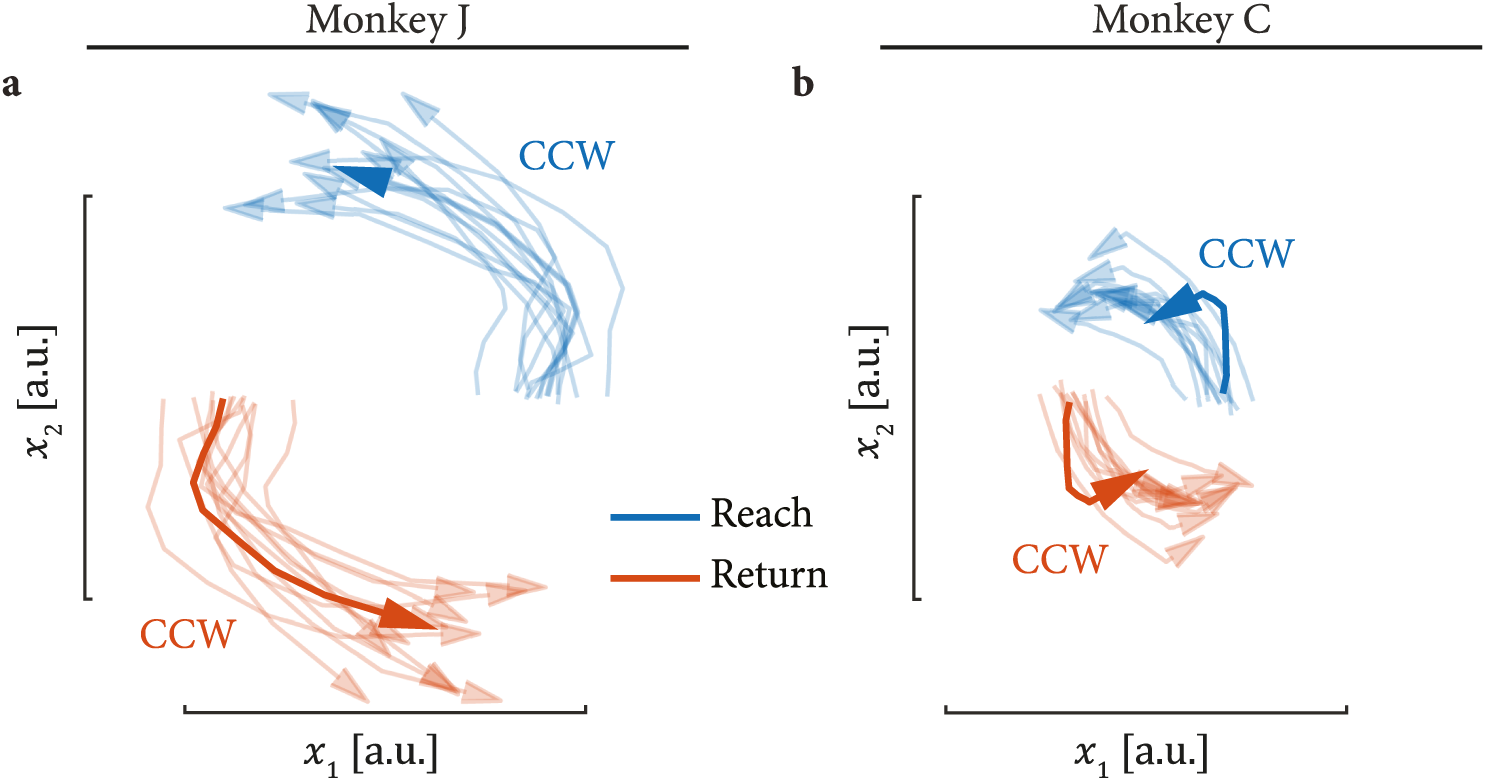
Similar to NDM, jPCA extracts rotations that are in the same direction during reach and return epochs. Notation is the same as in Fig. 4 for projections to 2D space extracted using jPCA.

**Supplementary Figure 8.**
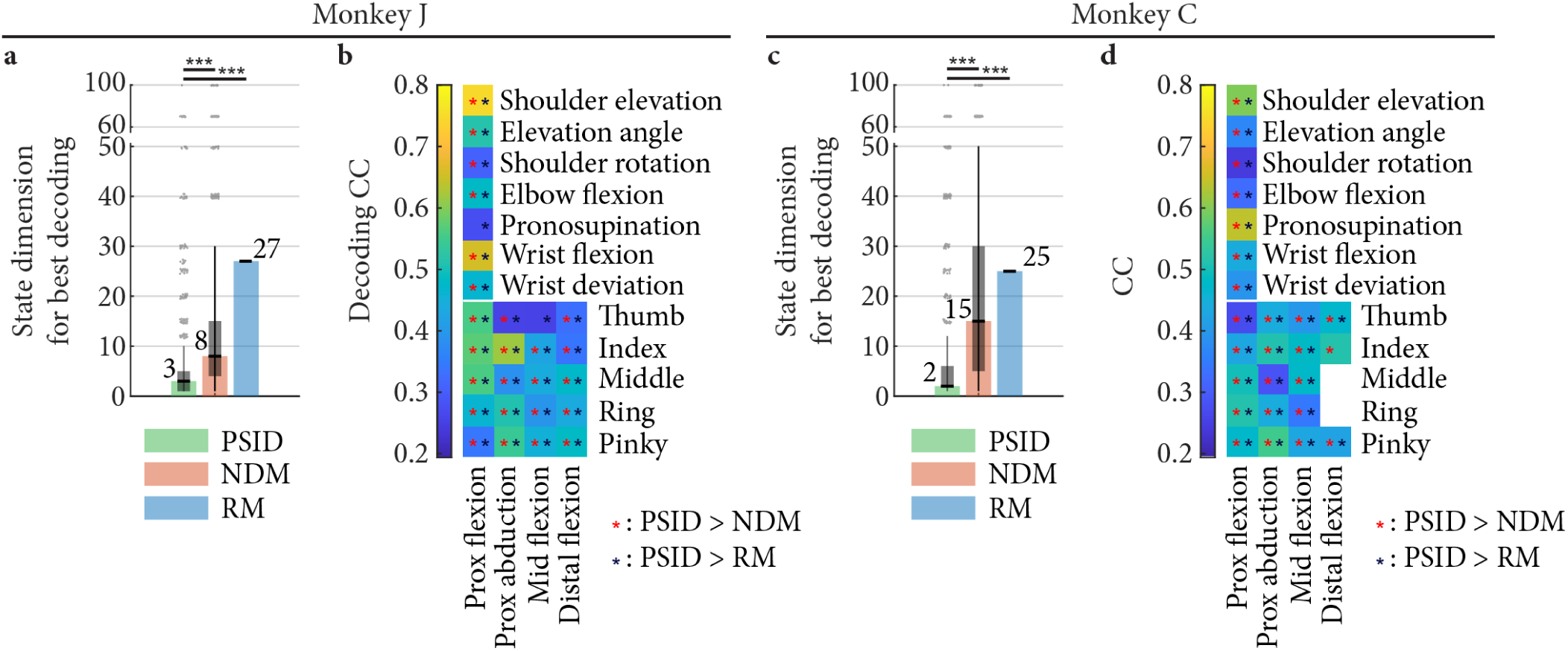
The PSID-extracted latent states with markedly lower dimension achieve significantly better decoding of almost all arm and finger joints. **(a)** The state dimension used by each method to achieve the best decoding for individual joints. For all methods, models are fitted to all joints as in Fig. 3. For PSID and NDM, models are fitted using various state dimensions; then for each joint, the latent state dimension is chosen to be the smallest value for which the decoding CC reaches within 1 s.e.m. of the best decoding CC possible for that joint among all latent state dimensions. Bars, boxes and asterisks are defined as in Fig. 3b. For better visualization of outliers, the vertical axis is broken. **(b)** Cross-validated correlation coefficient (CC) between the decoded and true joint angles is shown for PSID. Asterisks mark joints for which PSID results in significantly (P < 0.05) better decoding compared with NDM (red asterisk) or RM (dark blue asterisk). The latent state for each method is chosen as in (a). **(c)-(d)** Same as (a)-(b), for monkey C.

**Supplementary Video 1.**
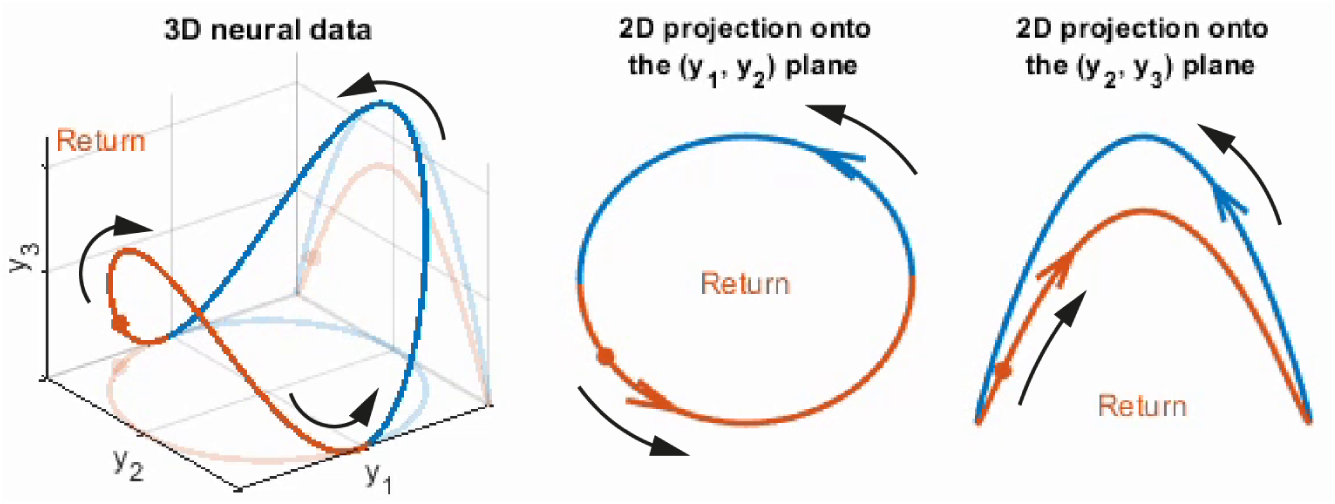
Visualization of how high-dimensional neural dynamics may contain 2D rotations both in the same and in opposite directions. The presented simulation depicts a hypothetical scenario where 3 dimensions of neural activity traverse a manifold in 3D space of which different projections reveal rotations in the same or opposite directions during reach vs return epochs. Among all projections, PSID can find the projection corresponding to the behaviorally relevant neural dynamics (e.g. here the (*y*_2_ − *y*_3_) plane, if behavior is best predicted using the activity in this plane) whereas the standard behavior-agnostic NDM methods may find other projections (e.g. the (*y*_1_ − *y*_2_) plane).

## Supplementary Notes

### Supplementary Note 1: The distinction between primary and secondary signals

To clarify the difference between the signals *y*_*k*_ and *z*_*k*_ in equation (2), it is worth noting that in the formulation of equation (2), *y*_*k*_ is taken as the primary signal in the sense that the latent state *x*_*k*_ describes the complete dynamics of *y*_*k*_ that also includes its shared dynamics with the secondary signal *z*_*k*_. The designation of the primary and secondary signals (e.g. taking *y*_*k*_ to be the neural activity and *z*_*k*_ to be the behavior or vice versa) is interchangeable as far as the shared dynamics of the two signals are of interest and the choice of the primary signal only determines which signal’s dynamics are fully described beyond the shared dynamics. In this work we take the primary signal *y*_*k*_ to be the neural activity and the secondary signal *z*_*k*_ to be the behavior. This is motivated by the typical scenario in neuroscience and neuroengineering where the neural activity is often considered the primary signal and the goal is to learn how behavior is encoded in it or to decode behavior from it.

The term 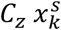 in equation (2), which we refer to as

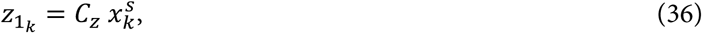

represents the part of the secondary signal *z*_*k*_ that is contributed by 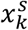 and thus shared with the primary signal. Any additional dynamics of the secondary signal that are not shared with the primary signal are modeled as the general independent signal *ϵ*_*k*_. If modeling these dynamics of the secondary signal is also of interest, after learning the parameters of equation (2), one could use the model to estimate 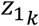 (Supplementary Note 4) and thus *ϵ*_*k*_ (as 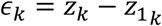) in the training data and then use standard dynamic modeling techniques (e.g. SID) to characterize the dynamics of *ϵ*_*k*_ in terms of another latent state-space model. But since these dynamics are independent of *y*_*k*_, such characterization would not be helpful in describing the encoding of *z*_*k*_ in *y*_*k*_ or in decoding of *z*_*k*_ from *y*_*k*_ and thus we will not discuss their identification, and only discuss their generation in our numerical simulations (Supplementary Note 7).

### Supplementary Note 2: Equivalent sets of parameters that can fully describe the model

We define 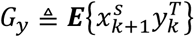 specifying the cross-covariance of *y*_*k*_ with the state at the next time step, 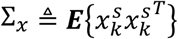 specifying the covariance of 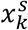 and 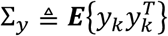 specifying the covariance of *y*_*k*_. From equation (2), it is straight forward to show that these covariances are related to the model noise statistics (equation (3)) via

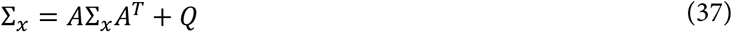

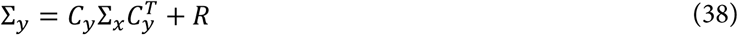

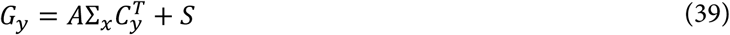

where equation (37) is also known as the Lyapunov equation^33,51^. The Lyapunov equation (37) has a unique solution for Σ_*x*_ if *A* is stable (i.e. the absolute value of all its eigenvalues are less than 1)^51^. For stable systems (models with a stable *A*), it is clear from equations (37)-(39) that there is a one to one relation between the set of parameters *A, C*_*y*_, *C*_*z*_, *G*_*y*_, Σ_*y*_, Σ_*x*_ and the set *A, C*_*y*_, *C*_*z*_, *Q, R, S*, and thus both sets can be used to describe the model in equation (2).

Equation (2) is known as the forward stochastic formulation for a linear state-space model. Given that only *y*_*k*_ and *z*_*k*_ are measurable real quantities and that the stochastic latent state 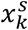 is not directly accessible, equation (2) is called an *internal* description for the signals *y*_*k*_ and *z*_*k*_^51^. This internal description is not unique and a family of infinitely many models with different 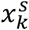 can describe the same *y*_*k*_ and *z*_*k*_. For example, any non-singular matrix *T*^′^ can transform equation (2) to an equivalent model with 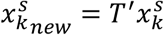, a process known as a similarity transform (or a change of basis). Moreover, Faurre’s stochastic realization problem shows that even beyond similarity transforms, there are non-unique sets of noise statistics (*Q, R*, and *S*) that give the exact same second order statistics for *y*_*k*_ ^33,51^. The unique and complete *external* description for *y*_*k*_ and *z*_*k*_ consists of their second order statistics. Thus, in the model learning problem, all models that give the correct external description are equally valid solutions. The set of parameters *A, C*_*y*_, *C*_*z*_, *G*_*y*_, Σ_*y*_, Σ_*x*_ are thus more suitable (compared with the equivalent set of parameters *A, C*_*y*_, *C*_*z*_, *Q, R, S*) for evaluating model learning because among this set, all parameters other than Σ_*x*_ are uniquely determined from second order statistics of *y*_*k*_ and *z*_*k*_, up to within a similarity transform^33,51^.

### Supplementary Note 3: Equivalent model formulation with behaviorally relevant states separated from the other states giving rise to equation (4)

Given the second order statistics of *y*_*k*_ (its auto-covariances at all possible time differences, see equation (51)), any set of parameters for equation (2) that would describe how the same second order statistics could be generated from a latent state 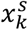 is known as a realization for *y*_*k*_ ^51^. We can rewrite equation (2) in an equivalent realization in which the behaviorally relevant states are clearly separated from the others. To do this, without loss of generality, we first assume that equation (2) is written as a minimal realization of *y*_*k*_, defined as a realization with the smallest possible state dimension *n*_*x*_^51^. For such a minimal realization, it can be shown that the pair (*A, C*_*y*_) is observable and the pair (*A, G*_*y*_) is reachable (Theorem 3.12 from ref. 51). Equivalently, both the neural observability matrix

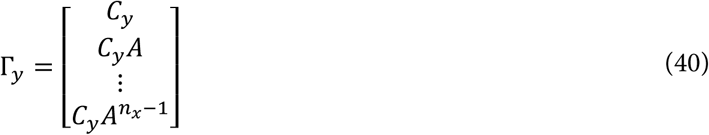

and the neural reachability matrix

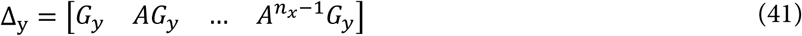

are full rank with rank of *n*_*x*_ (Theorems 3.4 and 3.7 from ref. 51).

Since not all latent states that contribute to the neural activity are expected to also contribute to a specific behavior of interest (equations (2) and (36)), the pair (*A, C*_*z*_) is not necessarily observable (i.e. it may not be possible to uniquely infer the full latent state 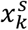 only from behavioral observations *z*_*k*_). In other words, the behavior observability matrix

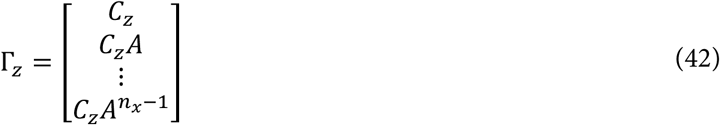

may not be full rank. We define *n*_1_ = rank(Γ_*z*_) as the number of latent states that drive behavior because as we show next, the latent state 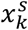 can be separated into two parts in a way that only *n*_1_ dimensions contribute to the behavior *z*_*k*_. We can show, by applying Theorem 3.6 from ref. 51 to the first and third rows of equation (2), that if *n*_1_ < *n*_*x*_, there exists a nonsingular matrix *T*^′^ that via the similarity transform

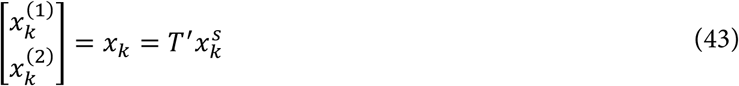

gives equation (4) as an equivalent formulation for equation (2).

### Supplementary Note 4: Kalman filtering and the equivalent forward innovation formulation

Given the linear state-space formulation of equation (2), it can be shown that the best prediction of *y*_*k*+1_ using *y*_1_ to *y*_*k*_ (denoted as *ŷ*_*k*+1|*k*_) in the sense of having the minimum mean-square error, and similarly the best prediction of *z*_*k*+1_ using *y*_1_ to *y*_*k*_ (denoted as 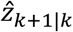) are obtained with the well-known recursive Kalman filter^51^, which can be written as

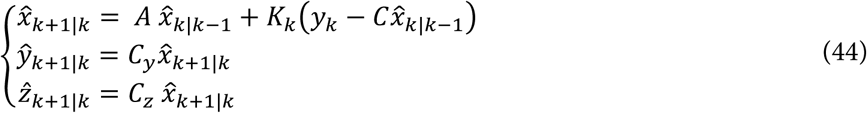

where the recursion is initialized with 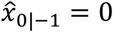 and *k*_*k*_ is the Kalman gain^51^ equal to

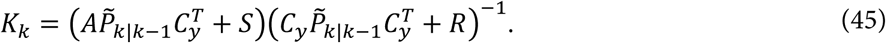

Here 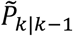 is the covariance of the error for one-step-ahead prediction of the state (i.e. covariance of 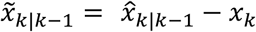 and can be computed via the recursive Riccati equation

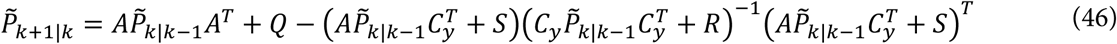

with the recursion initialized with *P*_0|−1_ = *R*_*y*_. The steady-state solution of Riccati equation can be obtained by replacing 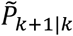 with 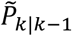 in the equation and solving for 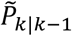. We will drop the subscript and denote the steady-state solution of equation (46) as 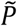 and the associated steady-state Kalman gain as *K*, which is obtained by substituting 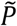 in equation (45).

Writing the outputs in terms of the Kalman filter states gives an alternative formulation for equation (2), which is known as the forward innovation formulation and is more convenient for deriving PSID. In particular, this formulation shows that the optimal estimate of the latent state is a linear function of the past neural activity. Based on this idea and the fact mentioned earlier that the best prediction of behavior and neural activity using past neural activity is a linear function of the latent state (equation (44)), we can show that linear projections of behavior and neural activity onto the past neural activity can be used to directly estimate the latent states from the data first, and then use the estimated latent states to learn the model parameters (Supplementary Note 5). The forward innovation formulation given by

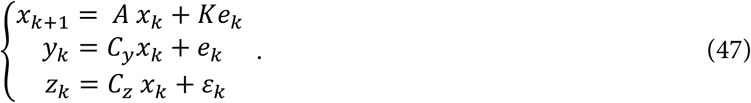

Here 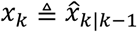, *k* is the steady-state Kalman gain and *e*_*k*_ is the innovation process, which is the part of *y*_*k*_ that is not predictable from its past values^33,51^. Equations (2) and (47) have different state and noise time-series but are equivalent alternative internal descriptions for the same *y*_*k*_ and *z*_*k*_ (Supplementary Note 2). The forward innovation formulation in equation (47) is more convenient (compared with the forward stochastic formulation in equation (2)) for the derivation of PSID. Specifically, by recursively substituting the previous iteration of equation (47) into its current iteration, it can be shown that

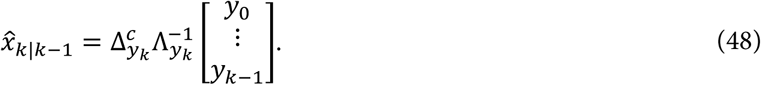

where

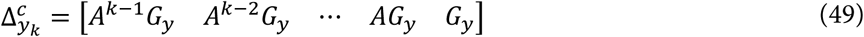

and

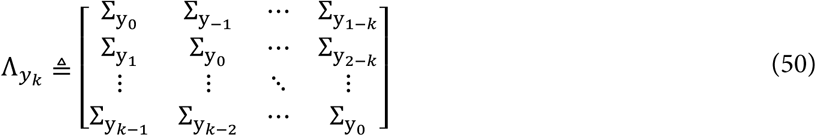

with the notation 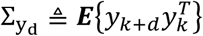 (Theorem 6 from ref. 33). This formulation reveals a key observation that enables identification of model parameters via a direct estimation of the latent state: the latent state in equation (47) (which is an equivalent formulation for equation (2)), is a linear function of the past *y*_*k*_. Moreover, from equation (2), it can be shown that for d ≥ 1

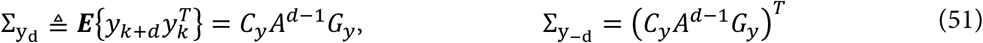

indicating that 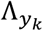 in equation (50) and thus the linear prediction function 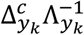 in (48) only depend on Σ_*y*_, *A, C*_*y*_ and *G*_*y*_^33,51^. Thus, from equations (44) and (48) it is clear that the only parameters that are needed for optimal prediction of *y*_*k*_ and *z*_*k*_ using past *y*_*k*_ are *A, C*_*y*_, *C*_*z*_, *G*_*y*_ and Σ_*y*_, which are all parameters that are uniquely identifiable within a similarity transform^33,51^ (Supplementary Note 2). As we confirm with numerical simulations, all these parameters can be accurately estimated using PSID (Supplementary Fig. 1).

### Supplementary Note 5: Derivations of PSID

#### PSID, stage 1: Extracting behaviorally relevant latent states

The central idea in PSID is that based on equations (44) and (48), the part of *z*_*k*_ that is predictable from past *y*_*k*_ is a linear combination of the past *y*_*k*_ and thus must lie in a subspace of the space spanned by the past *y*_*k*_. We use an orthogonal projection from future *z*_*k*_ onto past *y*_*k*_ to extract the part of *z*_*k*_ that is predictable from past *y*_*k*_, which leads to the direct extraction of the behaviorally relevant latent states from the neural and behavior data *y*_*k*_ and *z*_*k*_, even before the model parameters are known. Given the extracted latent states, the model parameters can then be estimated using least squares based on equation (4).

In the first stage of PSID, the part of *z*_*k*_ that is predictable from past *y*_*k*_ is extracted from the training data by projecting the future *z*_*k*_ values onto their corresponding past *y*_*k*_ values. To find the projection, for each time *k*, we consider the corresponding ‘past’ and ‘future’ to be the previous *i* samples and the next *i* − 1 samples respectively, with *i* being a user specified parameter termed the projection horizon. For each sample *y*_*k*_ with *i* ≤ *k* ≤ *N* − *i*, the previous (past) *i* samples (from *y*_*k*−*i*_ to *y*_*k*−1_) are all stacked together as the (*k* − *i* + 1)th column of one large matrix 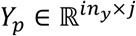 (with *j* = *N* − 2*i* + 1); correspondingly, for each sample *y*_*k*_ with *i* ≤ *k* ≤ *N* − *i*, that sample together with the next (future) *i* − 1 samples (from *y*_*k*_, to *y*_*k*+*i*−1_) are all stacked together as the (*k* − *i* + 1)th column of one large matrix 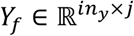 (equation (5)). Analogously, we form matrices 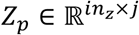 and 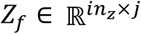 from *z*_*k*_ (equation (6)). Thus, *Z*_*f*_ and *Y*_*p*_ have the same number of columns with each column of *Z*_*f*_ containing some consecutive samples of behavior while the corresponding column in *Y*_*p*_ contains the previous *i* samples from neural activity. The goal is to find the part of *Z*_*f*_ that is linearly predictable from corresponding columns of *Y*_*p*_ (i.e. the behavior in each column of *Z*_*f*_ from its past neural activity). The linear least squares solution for this prediction problem has the closed form solution given in equation (7)^33,51^, which is in the form of a projection from future behavior onto past neural activity. We show below that this projection can be decomposed into the multiplication of an observability matrix for behavior and a running estimate of the Kalman estimated latent states, which will then enable the estimation of model parameters using the estimated latent states.

First, note that the least squares solution of equation (7) can also be written as

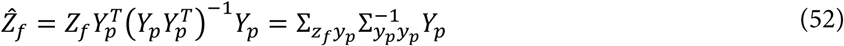

where 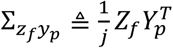 and 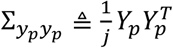 are sample covariance matrices for the covariance of past neural activity with future behavior and past neural activity, respectively, computed using their observed time-samples from equations (5) and (6). Sample covariance estimates are asymptotically unbiased and thus for *j* → ∞ they would converge to their true value in the model^33,51^. Consequently, for the model in equation (2), it can be shown (by replacing samples covariances with exact covariances from the model) that for *j* → ∞, 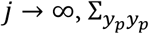 converges to 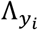 defined per equation (50) and 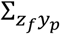 converges to

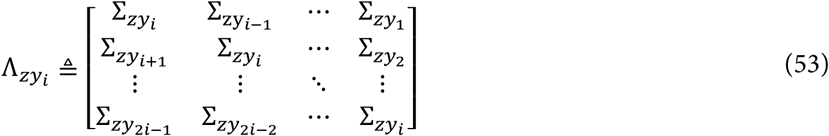

where we are using the notation 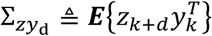. From equation (2) it can be shown that

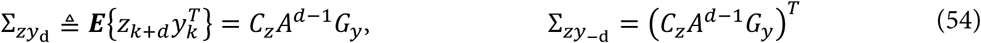

which has a form similar to equation (51). Substituting into the definition of 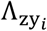 from equation (53) gives

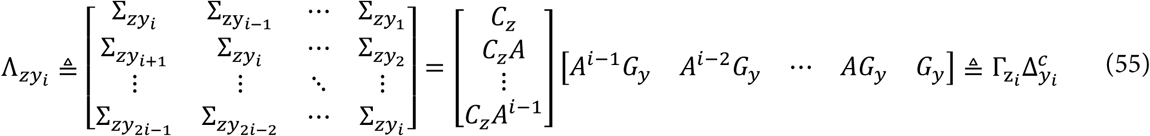

where 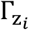 is termed the extended observability matrix for the pair (*A, C*_*z*_) and 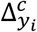 is termed the reversed extended controllability matrix for the pair (*A, G*_*y*_)^33^. Moreover, based on equation (48), the Kalman filter prediction at time *k* using only the last *i* observations (*y*_*k*−*i*_ to *y*_*k*−1_) can be written in terms of 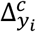 (equation (55)) as

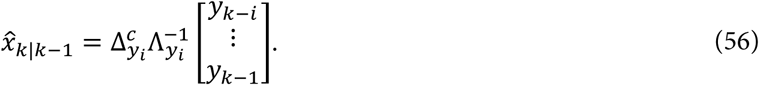

Thus, for *j* → ∞, equation (52) can be written as

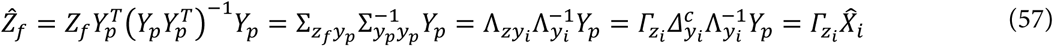

where columns of 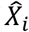 are Kalman estimates obtained using the past *i* observations of *y*_*k*_ (from equation (56)). Before we use equation (57) to conclude the derivation of the first stage of PSID, it is useful for the derivation of the second stage to note that if we repeat the above steps for the projection of *Y*_*f*_ onto *Y*_*p*_, we will get

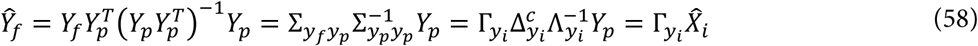

where 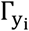 is the extended observability matrix for the pair (*A, C*_*y*_) and 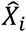 are *the exact same* Kalman states as in equation (57).

Equation (57) shows how 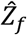, which is the projection of future behavior onto past neural activity and is directly computable from data, can be decomposed into the extended behavior observability matrix 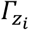 and the Kalman states 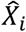. This decomposition allows us to estimate the latent states even before the model parameters are learned and paves the way for subsequent learning of the model parameters. The decomposition can be performed by taking singular value decomposition (SVD) from equation (57) (shown in equation (9)), which gives:

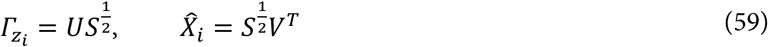

Note that the above is only one of many valid decompositions since multiplying any non-singular matrix *T* onto 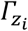 from the right and its inverse *T*^−1^ onto 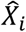 from the left amounts to a similarity transform and gives an equivalent model with a different basis^33^. Without loss of generality, we assume that the latent states are not trivial linear combinations of each other and thus 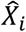 is full rank. Given that only *n*_1_ states drive behavior (Supplementary Note 3), 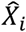 as well as 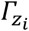 will have rank of *n*_1_. Indeed, *n*_1_ was defined as the rank of the behavior observability matrix Γ_*z*_ and for a sufficiently large horizon *i* (i.e. *i* ≥ *n*_1_ is sufficient but not necessary), the rank of the extended behavior observability matrix 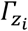 will also be *n*_1_^51^. For *j* → ∞, as shown earlier, equation (57) holds exactly and thus the row rank of 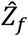 and the number of its non-zero singular values will be equal to the rank of 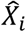 and 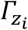, which is *n*_1_ (Supplementary Note 3). For finite data (*j* < ∞), it is expected that an approximation of this relation will hold and thus one could find *n*_1_ via inspection of the singular values of 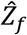. In Methods, we instead proposed a more general method of using cross validation to find both *n*_1_ and *n*_*x*_, which doesn’t require an ad-hoc determination of which singular values are notably larger than the others. Regardless of how *n*_1_ is determined, keeping the top *n*_1_ singular values from the SVD, we can extract 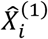 as in equation (13). Note that in this stage, keeping of the top singular values ensures that the states that describe the behavior best (i.e. best approximate 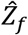) are prioritized.

Having decomposed 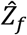 into 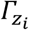 and 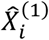, determining the model parameters from these matrices is straight forward and there are multiple possible ways to accomplish this. We take an approach in the spirit of stochastic algorithm 3 from ref. 33 in SID, and use the state matrix 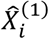 to estimate the model parameters. This method has the advantage of guaranteeing that the estimated noise statistics are positive semi-definite, which is necessary for the model to be physically meaningful^33^. We first compute the subspace for the latent states at the next time step (having observed *Y*_*i*_ as defined in equation (5) in addition to *Y*_*p*_, i.e. having observed the past *i* + 1 samples) by projecting 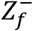 onto 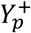 (equation (8)). Similar to equation (57), this projection can be decomposed as

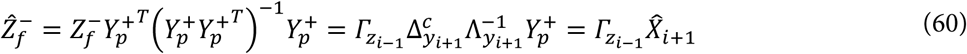

where 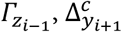 and 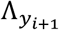 are defined similar to equations (55) and (50) and columns of 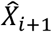 are Kalman estimates obtained using the past *i* + 1 observations of *y*_*k*_ (from equation (56)). From the definition of observability matrix, it is clear that 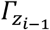 can be computed by removing the last block row of 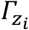 (equation (12)). 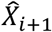 can then be computed (in the same basis as 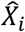) by multiplying both sides of equation (60) with 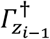 from the left (equation (13)). We then take columns of 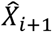 and 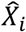 as samples of the current state and the corresponding next state (i.e. 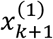 and 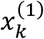 from equation (4)) respectively, and based on equation (4), compute the least squares estimate for *A*_11_ that is given in equation (14). This concludes the extraction of behaviorally relevant latent states and the estimation of the segment of the state transition matrix *A* that is associated with these states (i.e. *A*_11_). In the next stage of PSID, we extract the behaviorally irrelevant latent states (optional) and estimate the rest of the *A* matrix and all other model parameters using the extracted states to conclude the full derivation.

#### PSID, stage 2: extracting behaviorally irrelevant latent states

So far we have extracted the behaviorally relevant latent states as the key first step toward learning the model parameters. To find any remaining behaviorally irrelevant states, we need to find the variations in neural activity that are not explained by the behaviorally relevant latent states. We thus first remove any variations in *Y*_*f*_ (and 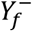) that lies in the subspace spanned by the extracted behaviorally relevant states 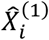 (and 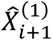) (*i* is horizon as defined previously), and then apply a procedure akin to the standard SID to the residual. The least squares solution for the best linear prediction of *Y*_*f*_ using 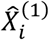 is given by equation (15), and is termed 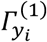. This solution can be thought of as the neural observability matrix associated with the behaviorally relevant states 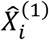 (equation (58)). Thus, similar to equation (60), the associated observability matrix for 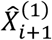 can be computed by removing the last block row from the solution (equation (17)). We then subtract the best prediction of 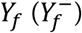 using 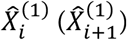 from it as shown in equation (16) (equation (18)), and call the result 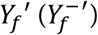. In other words, 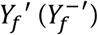 is the part of 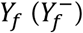 that does not lie in the space spanned by 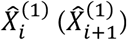. Given that 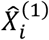 and thus 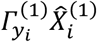 (i.e. the linear prediction of *Y*_*f*_ using 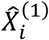) are of rank *n*_1_ and that *Ŷ*_*f*_ (i.e. the projection of *Y*_*f*_ onto *Y*_*p*_) is of rank *n*_*x*_ (equation (58)), the projection of 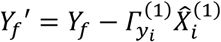 (i.e. residual future neural activity) onto *Y*_*p*_ will be of rank *n*_2_ = *n*_*x*_ − *n*_1_. A similar procedure to what was applied to *Z*_*f*_ (and 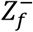) to find 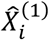 (and 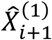) can be applied to *Y*_*f*_′ (and 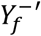) to extract the *n*_2_ remaining states 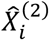 (and 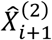) (steps 11-14 from Table 1). Of note is that in this stage, keeping the top singular values after SVD (equation (21)) ensures that the remaining states that describe the unexplained neural activity best (i.e. best approximate 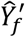) are prioritized.

The above concludes the extraction of behaviorally irrelevant latent states. Concatenating the states extracted from both stages (i.e. 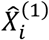 and 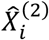 as well as 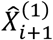 and 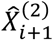) together as in equation (26) concludes the extraction of all latent states, including behaviorally relevant and irrelevant ones. Given the fully extracted latent states, we then follow a similar approach as was taken before for *A*_11_ (equation (14)), to find the least squares estimate for *A*_12_ and *A*_22_ (equation (27)), *C*_*y*_ (equation (29)) and *C*_*z*_ (equation (30)). Finally, the residuals from the least squares solutions to equations (14), (27) and (29) provide estimated values for *w*_*k*_ and *v*_*k*_ at each time step and thus we compute the sample covariance of these residuals to find the noise covariance parameters (equation (32)). This concludes the estimation of all model parameters.

Finally, in addition to equation (30), another viable alternative for finding the parameter *C*_*z*_ is to learn it using linear regression, which is the procedure needed for the standard SID to relate its extracted latent state to behavior and we use in this paper for both SID and PSID. Since *C*_*z*_ is not involved in the Kalman filter recursions (first 2 rows of equation (44)), it does not have any effect on the estimation of latent states from *y*_*k*_ and it only affects the later prediction of *z*_*k*_ from those latent states. Consequently, we can use the other identified parameters to apply Kalman filter to the training *y*_*k*_ and estimate the latent states 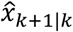 (equation (44)). We can then use linear regression to find the *C*_*z*_ that minimizes the prediction of *z*_*k*_ using 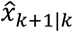 as

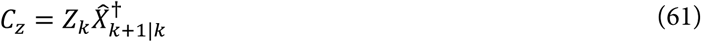

where columns of *Z*_*k*_ contain *z*_*k*_ for different time steps and columns of 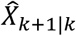 contain the corresponding 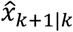 estimates for those time steps. The advantage of using this alternative estimation of *C*_*z*_ is that 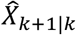 (used in equation (61)) are more accurate estimates of the latent states obtained using all past observations whereas 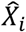 (used in equation (30)) are less accurate estimates obtained using only the past *i* observations.

### Supplementary note 6: Standard SID as a special case of PSID and the asymptotic characteristics of PSID

As a review of the standard SID, we refer the reader to chapter 8 from ref. 51 and chapter 3 from ref. 33. For *n*_1_ = 0, PSID (Table 1) reduces to the standard SID (specifically to stochastic algorithm 3 from ref. 33). This is because if *n*_1_ = 0, no behaviorally relevant states 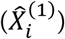 will be extracted leaving all variation of *Y*_*f*_ to be identified in stage 2 of PSID, which is similar to using standard SID. Thus, the extracted 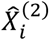 in this case will be the same as the 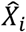 that is obtained from applying SID on *y*_*k*_. So to compare PSID with SID, we simply use PSID with *n*_1_ = 0.

As a generalization of the abovementioned version of SID (i.e. stochastic algorithm 3 from ref. 33), PSID has similar asymptotic characteristics. In some other variations of SID (for example in stochastic algorithm 2 from ref. 33 and in Algorithm A in section 8.7 from ref. 51), instead of applying SVD to *Ŷ*_*f*_, SVD is applied to the empirical cross-covariance 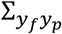 to decompose it into 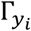 and 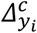 (equation (58)), giving an estimation of these matrices which for *j* → ∞ is unbiased^33^. From this decomposition, model parameters *A, C*_*y*_, and *G*_*y*_ can then be—extracted—*C*_*y*_ as the first block of 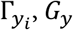, as the last block of 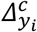, and *A* with a least squares solution within blocks of 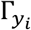 (for details see the SID variants mentioned in the previous sentence). However, this approach cannot guarantee that for finite data (*j* < ∞) the identified *A, C*_*y*_, and *G*_*y*_ will be associated with a positive real covariance sequence (i.e. Faurre’s stochastic realization may have no solution)^33^. In the alternative approach taken by PSID (and its special case, stochastic algorithms 3 from ref. 33), *A* and *C*_*y*_ are computed as least squares solution of forming equation (2) with 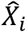 taken as the value of the latent state and *G*_*y*_ is identified later based on the residuals of the least squares solution. This approach cannot guarantee an asymptotically unbiased estimate of *G*_*y*_ (unless *i* → ∞ in which case Kalman estimates in equation (56) will be exact), but it guarantees that even for finite data (*j* < ∞) the identified parameters will be associated with a positive real covariance sequence^33^, which is essential for the model to be physically meaningful^33^.

### Supplementary Note 7: Generating random model parameters for simulations

For a model with given *n*_*x*_ and *n*_1_, *A* was generated by first randomly generating its eigenvalues and then generating a block diagonal real matrix with the randomly selected eigenvalues (using MATLAB’s cdf2rdf command). We drew the eigenvalues with uniform probability from across the complex unit circle and then randomly selected *n*_1_ of the *n*_*x*_ to be later used as behaviorally relevant eigenvalues. As a technical detail, in both the original random generation of eigenvalues and in selecting *n*_1_ of them for behavior we made sure eigenvalues are either real valued or are in complex-conjugate pairs (as needed for models with real observations). To do this, we first drew 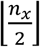 points with uniform probability from across the complex unit circle and then added the complex conjugate of each to the set of eigenvalues. If *n*_*x*_ was odd, we then drew an additional eigen value from the unit circle and set its angle to 0 or *π*, whichever was closer. Finally, to randomly select *n*_1_ of the *n*_*x*_ eigenvalues to be used as behaviorally relevant, we repeatedly permuted the values until the first *n*_1_ eigenvalues also formed a set of complex conjugate pairs or real values.

Next, we generated 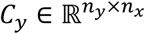 by drawing each element from standard normal distribution. We generated 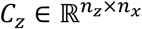 by drawing values from the standard normal distribution for the elements associated with the behaviorally relevant eigenvalues of *A* (or equivalently for the dimensions of *x*_*k*_ that drive behavior) and setting the other elements to 0 (see equation (4)).

For noise statistics *Q, R*, and *S*, we generated general random covariance matrices and applied random scaling factors to them to get a wide range of relative variances for the state noise *w*_*k*_ and observation noise *v*_*k*_. To do this, we first generated a random square matrix Ω of the size *n*_*x*_ + *n*_*y*_ by drawing elements from standard normal distribution and computed *L* = ΩΩ^*T*^, which is guaranteed to be symmetric and positive semi-definite. We next selected random scaling factors for the state noise *w*_*k*_ and the observation noise *v*_*k*_ by independently selecting two real numbers *a*_1_, *a*_2_ with uniform probability from the range (−1, +1). We then applied the following scaling to matrix *L* to get the noise statistics as

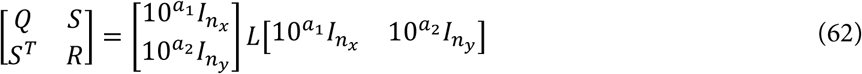

where *I*_*n*_ denotes the identity matrix of the size *n*. This is equivalent to scaling *v*_*k*_ by 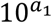 and independently scaling *w*_*k*_ by 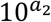 and generates a wide range of state and observation noise statistics.

Finally, to build a model for generating the independent behavior residual dynamics *ϵ*_*k*_ (which can be a general colored signal and is not assumed to be white), we generate another random dynamic linear SSM with independently selected latent state dimension of 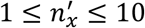 and parameters generated as explained above for the main model. We will refer to this model as the behavior residual dynamics model. To diversify the ratio of behavior dynamics that are shared with neural activity (equation (36)) to the residual behavior dynamics (i.e. *ϵ*_*k*_), we draw a random number *α*_3_ in the range (0, +2). We then multiply the rows of the *C*_*z*_ parameter in the behavior residual dynamics model with different scalar values such that for each behavior dimension *m*, the shared-to-residual ratio, defined as the ratio of the std of the term 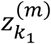 (equation (36)) to the std of the term 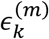, is equal to 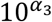.

